# Evolutionary modelling of HCV subtypes provides rationale for their different disease outcomes

**DOI:** 10.1101/2021.02.02.429470

**Authors:** Hang Zhang, Ahmed A. Quadeer, Matthew R. McKay

**Affiliations:** Department of Electronic and Computer Engineering, The Hong Kong University of Science and Technology, Clear Water Bay, Hong Kong, China; Department of Chemical and Biological Engineering, The Hong Kong University of Science and Technology, Clear Water Bay, Hong Kong, China

## Abstract

Hepatitis C virus (HCV) is a leading cause of liver-associated disease and liver cancer. Of the major HCV subtypes, patients infected with subtype 1b have been associated with having a higher risk of developing chronic infection, cirrhosis and hepatocellular carcinoma. However, underlying reasons for this increased disease severity remain unknown. Here, we provide an evolutionary rationale, based on a comparative study of fitness landscape and in-host evolutionary models of the envelope glycoprotein 2 (E2) of HCV subtypes 1a and 1b. Our analysis demonstrates that a higher chronicity rate of subtype 1b may be attributed to lower fitness constraints, enabling 1b viruses to more easily escape antibody responses. More generally, our results suggest that differences in evolutionary constraints between HCV subtypes may be an important factor in mediating distinct disease outcomes. Our analysis also identifies antibodies that appear to be escape-resistant against both subtypes 1a and 1b, providing directions for the design of HCV vaccines having cross-subtype protection.

## I. Introduction

HCV is a highly mutable single-stranded RNA virus [1] which is estimated to affect 2.5% of the global population [2]. Approximately 15% to 25% of infected people clear the virus spontaneously, while the remaining develop chronic liver disease, commonly leading to cirrhosis and hepatocellular carcinoma (HCC) [3]. Although therapeutic treatments for chronic HCV infection have improved since the introduction of direct-acting antivirals (DAAs), these drugs are only administered to a limited number of infected individuals because of high cost [4] and low diagnostic rate [5]. The effectiveness of DAAs also has limitations due to the inability to prevent reinfection [6] and the emergence of drug-resistant variants [7]. Thus, for complete eradication of HCV, developing an effective vaccine is essential.

HCV is classified into 7 major genotypes and 67 subtypes [8]. Among the major genotypes, genotype 1 is the most prevalent, representing roughly half of all HCV infections worldwide [2]. Of the genotype 1 infections for which subtypes have been specified, 99% are either subtype 1a or 1b [9], with these two subtypes having approximately equal prevalence worldwide [9]. In terms of disease outcome, subtype 1b is particularly problematic, with ~92% of patients exposed to this subtype having been found to develop chronic infection. This is more than double the corresponding percentage of patients exposed to 1a and other subtypes [10]. Moreover, subtype 1b infections have been reported to increase the risk of developing cirrhosis [11], and present almost double the risk of developing HCC compared to 1a and other subtypes [12]–[15].

Despite these observations, it remains unclear as to why subtype 1b infections may lead to more severe disease outcomes. This may be attributed to various reasons, including differences in viral phenotype, host-specific immune factors, demographics, etc. One plausible explanation is that subtype 1b may evolve under less rigid fitness constraints than other subtypes, allowing it to escape immune responses more easily and to more effectively propagate infection. Evidence of a similar phenomenon has been presented for HIV, for which the fitness of the infecting subtype has been reported to be a factor associated with disease outcome [16]. For HCV, it is not yet known whether different subtypes exhibit markedly different fitness properties, and if so, whether this could be a significant factor in determining disease outcome.

We aim to address these questions by using computational modelling, leveraging sequence data as well as available clinical and experimental data. Our approach is to first infer a model for the fitness landscape of HCV E2 (the primary target of neutralizing antibodies) for subtype 1b, which we validate against in-vitro fitness measurements, and compare it with an analogous model for subtype 1a that we had inferred previously [17]. Our comparative analysis suggests E2 1b to be more tolerant to mutations compared to E2 1a, indicating that it may be easier for subtype 1b viruses to evade immune responses. This is further corroborated by an analysis of the average time it takes for each subtype to escape from antibody responses, which we quantify by using population-genetics-based models of in-host viral evolution [17]. Our analysis, in general, points to significant differences in viral evolutionary constraints experienced by different subtypes of HCV, which may contribute to observed subtype-specific differences in disease outcomes. We additionally employ the evolutionary models to identify potentially escape-resistant human monoclonal antibodies (HmAbs) against subtypes 1a and 1b, which may assist in the rational design of a vaccine that is effective against the most prevalent HCV subtypes worldwide.

## II. Results

### A. Inference and validation of the HCV E2 1b fitness landscape

We inferred a computational model for the fitness landscape of E2 1b using the sequence data available at the HCV-GLUE database [18], [19] (see Methods for details). This involved obtaining a maximum entropy (least-biased) probabilistic model—a “prevalence landscape” that captures the probability of observing a virus with a particular E2 protein sequence in circulation. In this model, the probability of any sequence x = [*x*_1_ *x*_2_,…,*x_N_*] can be expressed as

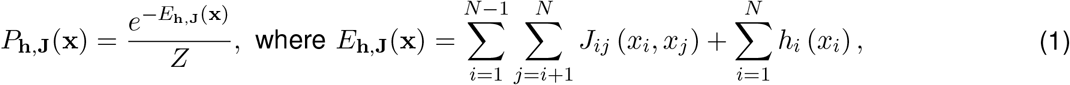

where h is the set of all fields that represent the effect of mutations at a single residue, J is the set of all couplings that represent the effect of interactions between mutations at two different residues, and *Z* = ∑_x_ *e*^-*E*_h,J_(x)^ is a normalization factor. The quantity *E*_h,j_(x) represents the energy of sequence x, which is inversely related to its prevalence. Inferring a maximum entropy model involves choosing the model parameters (fields and couplings) such that the single and double mutant probabilities obtained from the model match with those of the multiple sequence alignment (MSA). Maximum entropy models have been used for inferring the fitness landscape of HCV polymerase protein [20] and of several proteins of HIV [21]–[25], and for designing a T cell based HIV vaccine candidate, shown to be immunogenic in rhesus macaques [26]. The maximum entropy model inference for E2 protein is challenging as it involves estimating a large number of model parameters, a consequence of the high mutational diversity of E2 compared to other HCV proteins (Supplementary Fig. S1). To tackle this problem, we used an efficient computational approach introduced in [25] to infer fitness landscapes of the HIV envelope protein. The method was also applied to infer a fitness landscape for the E2 protein of HCV subtype 1a [17].

The single and double mutant probabilities obtained from our inferred model for HCV E2 subtype 1b aligned well with those of the MSA (Supplementary Fig. S2a, b). Other statistics, such as triple mutant probabilities and the distribution of the number of mutations computed from our inferred model, although not explicitly included while training, also matched well with those of the MSA (Supplementary Fig. S2c, d). These results indicate that our inferred prevalence landscape model accurately captures the statistical variations in the observed E2 1b sequence data.

Comparing fitness predictions from the inferred model with in-vitro infectivity measurements reported by four different studies [27]–[30] demonstrated that the inferred landscape for E2 1b is a reasonably good representative of the underlying protein fitness landscape (Fig. 1a). Specifically, a strong negative Spearman correlation (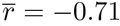; see Methods for details) was observed between the model-predicted energies (inversely related to prevalence; see Eq. (1)) and measured fitness values. This accuracy is commensurate with fitness predictions reported by studies of other proteins [17], [20]–[22], [24], [25].

**Fig. 1:**
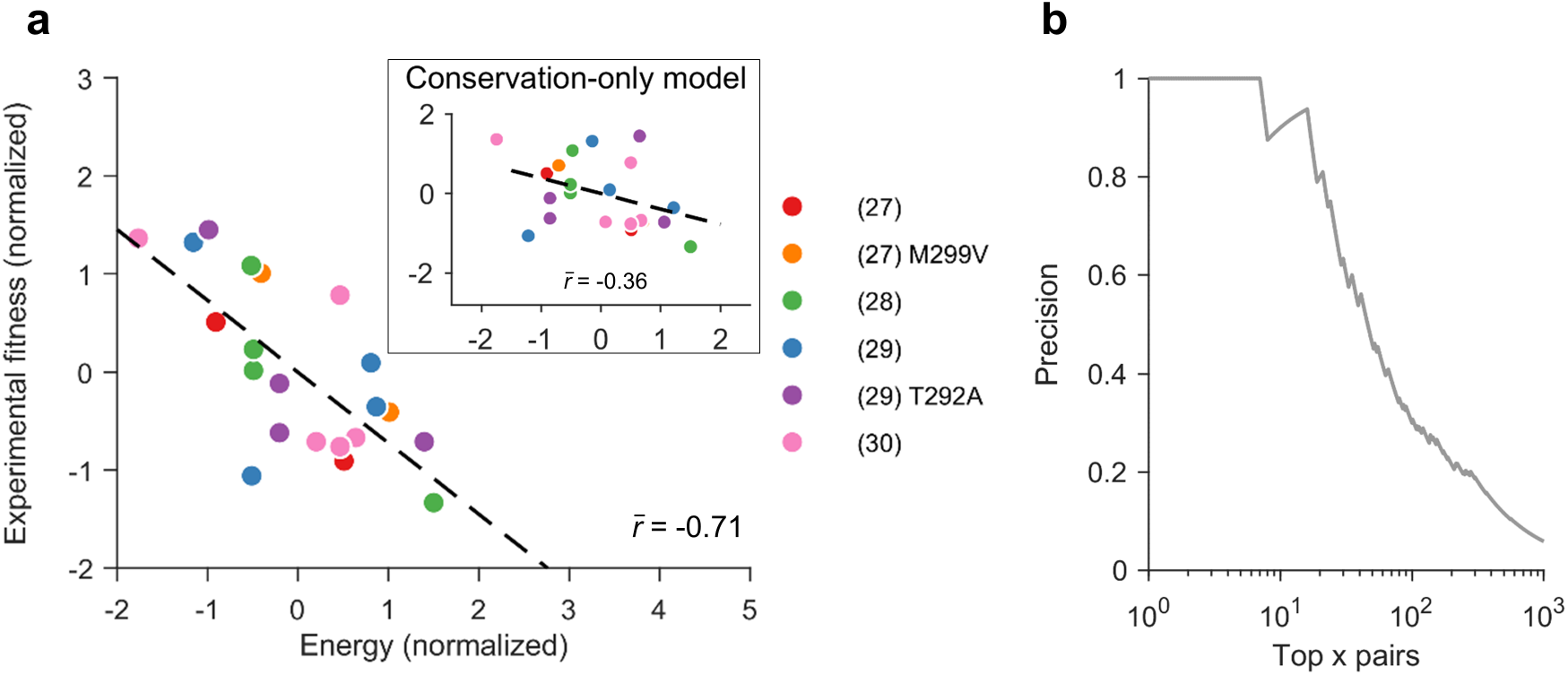
Validation of the inferred E21b fitness landscape. (**a**) Normalized experimental fitness measurements correlate strongly with the energy computed from the inferred landscape. In contrast, a much lower correlation is observed for the conservation-only model (Supplementary Text S1) that does not take couplings into account (inset). References for fitness (infectivity) measurements are shown in the legend. In [27] and [29], E2 fitness measurements were reported in two different E1 backgrounds. In [27], one background involved methionine (M) at residue 299, while the other involved valine (V). Similarly, in [29], one background involved threonine (T) at residue 292, while the other involved alanine (A). (**b**) Precision of contact predictions vs. the top x pairs according to the inferred model couplings. Precision is the proportion of top x pairs that are truly in contact. Two residues were assumed to be in contact if their carbon-alpha atoms were less than 8Å apart according to the available E2 1b crystal structure (PDB ID: 6MEI).

In addition to mutations at individual residues, interactions between mutations at different residues have been shown to be important contributors to HCV fitness [17], [31]. Thus, we compared the predictions of our model, which takes into account residue interactions via couplings, with those of a model based only on amino acid conservation (or single mutant probabilities; see Supplementary Text S1 for details). Compared to our model, the conservation-only model provided a much lower correlation 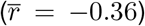 between the model-predicted energy and in-vitro infectivity measurements (Fig. 1a, inset), corroborating the importance of incorporating residue interactions in determining fitness. This suggests that the fitness effect of individual mutations in HCV also depends on the sequence background in which they are introduced. Moreover, the strong couplings of maximum entropy models are known to be informative of contacts in the protein tertiary structure [32]–[34]. The inferred E2 1b model couplings were also found to be good predictors of contacts in the E2 1b protein structure [PDB ID: 6MEI] (Fig. 1b; see Methods for details). The contact prediction from the model couplings was comparable with direct coupling analysis (DCA) [34], a standard contact prediction method (Supplementary Fig. S3).

### B. Fitness models indicate subtype 1b to be less evolutionarily constrained than subtype 1a

We compared the fitness landscape model of HCV E2 1b with an analogous model developed previously for E2 1a [17] to investigate whether the observed differences in disease outcome of the two subtypes’ infections may be related to the associated fitness constraints. We first compared the fitness landscapes based on two standard metrics used for quantifying landscape ruggedness: autocorrelation and neutrality [35]–[37]. Autocorrelation quantifies the average change in fitness as one moves randomly along the landscape (see Methods for details). We observed that the decay of autocorrelation of the E2 1b landscape was slower than that of 1a (Fig. 2a, left panel), suggesting that the average change in fitness while moving along the landscape of 1b is less than 1a. The neutrality associated with each landscape quantifies the maximum number of mutation steps one can take on average without much change in fitness while moving randomly along the landscape (see Methods for details). Our results showed that the E2 1b landscape was more neutral than 1a (Fig. 2a, right panel), further suggesting that the average fitness cost upon mutation in 1b is lower than 1a. These results were qualitatively robust to the choice of the involved parameters (Supplementary Fig. S4).

**Fig. 2:**
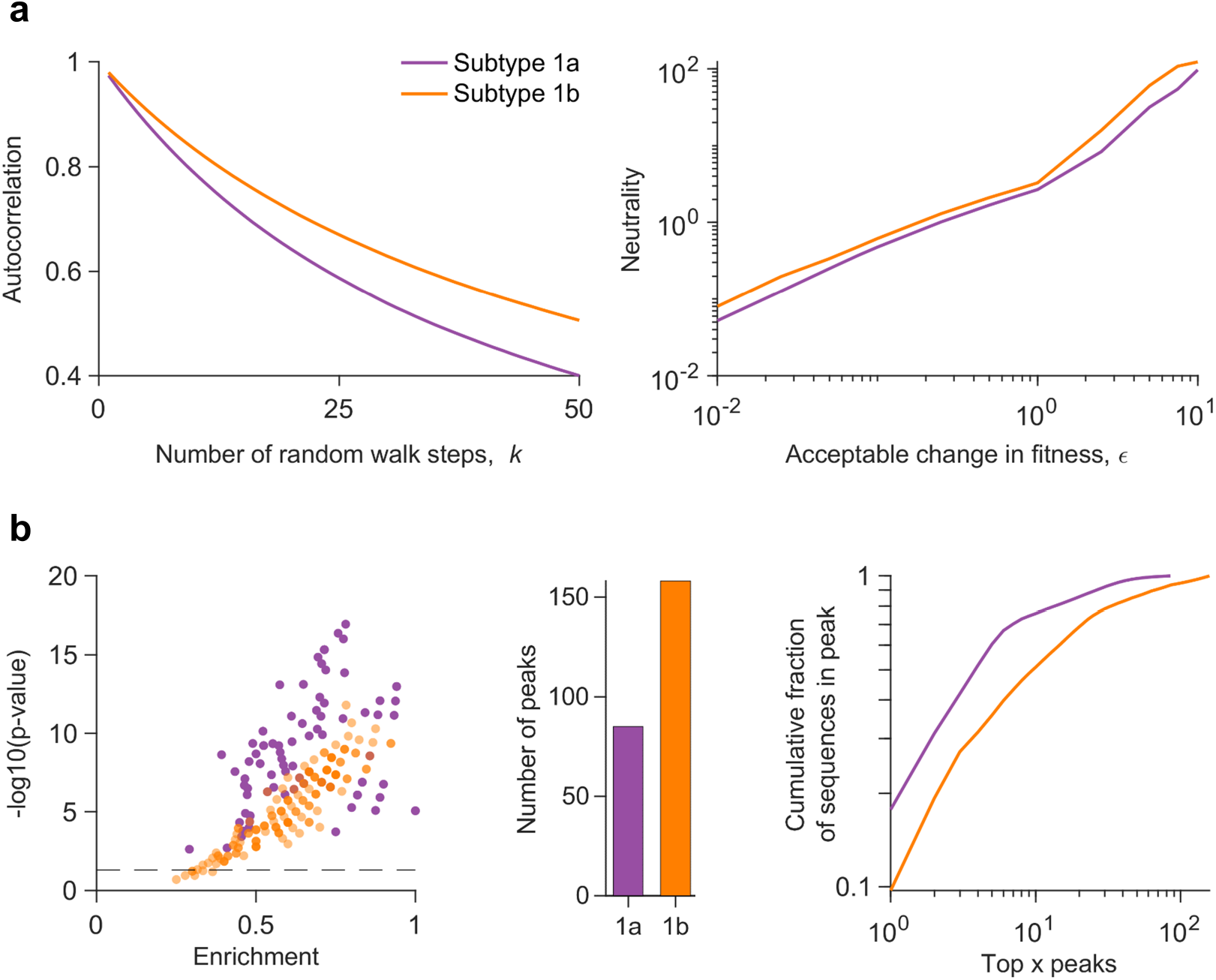
Comparison of the fitness landscapes of E2 1a and 1b. (**a**) (Left panel) Comparison of the autocorrelation of the sequence energies of each subtype’s landscape. The x-axis denotes the number of random walk steps (k) for which the autocorrelation was computed. The starting sequences were chosen within a Hamming distance (D) of 5 from the MSA (see Methods for details). (Right panel) Comparison of the neutrality of the landscape of each subtype. Neutrality was computed for L = 500 random walk steps. A random step in the walk was accepted only when the change in fitness from the sequence at the previous step was within a small value, e (shown on the x-axis) (see Methods for details). (**b**) (Left panel) Statistically significant enrichment of almost all peak sequences in known escape mutations (listed in Supplementary Table S1), for each subtype. Enrichment is the fraction of known escape mutations within all mutations in a peak sequence, and the p-value measures the probability of observing by random chance at least the observed number of escape mutations among all observed mutations in a peak sequence (see Methods for details). The dashed horizontal line is the cut-off for statistically significant results (p-value < 0.05), with all peak sequences above this line being classed as statistically significant. (Middle panel) Number of peaks observed in each landscape. (Right panel) The cumulative fraction of sequences associated with the peaks versus the top x peaks in each landscape, plotted on a log-log scale.

We also compared the two landscapes using a low-dimensional representation that characterises each landscape by its local “peaks”, representing fitness maxima. Such local peaks have been shown to be informative of immune escape pathways employed by HIV [38], and of the evolution of poliovirus under vaccine-induced and natural selective pressures [37]. Peaks were obtained by following a steepest ascent walk from each sequence in the MSA. In the previous HIV study [38], sequences representing each peak in the landscape, referred to as “peak sequences”, were found to be strongly enriched in HLA-associated mutations driven by host immune responses. Investigating the peak sequences of each subtype for HCV E2 revealed that almost all of the peak sequences of both subtypes were statistically significantly enriched in known escape mutations from E2-specific HmAbs [39]–[45] (Fig. 2b left panel; see Methods for details). This suggests that peaks in the fitness landscapes of E2 may be representative of immune escape pathways employed by HCV for both subtypes. This result was also qualitatively robust to the change of parameter values involved in the landscape inference (Supplementary Fig. S5). We observed that there were far more peaks in the landscape of 1b than 1a (Fig. 2b, middle panel), suggesting that more escape pathways may be available for subtype 1b than 1a. Rank-ordering the peaks according to the fraction of MSA sequences represented by each peak revealed that the same number of top peaks in 1b comprised a smaller fraction of sequences than 1a (Fig. 2b, right panel). This indicates that the sequences are more spread out across peaks in 1b, which is suggestive of relatively diverse escape pathways utilized by subtype 1b in comparison to 1a. These qualitative results were also robust to the change of parameter values involved in the landscape inference (Supplementary Fig. S6). We additionally confirmed, using a procedure similar to [37], [38], that the peaks observed in each landscape are not an artefact of finite sampling, but arise due to the interplay between mutations at different residues in each subtype (Supplementary Text S2).

Overall, this comparison of fitness landscapes demonstrates that intrinsic fitness constraints governing the evolution of HCV subtypes may differ significantly. Our results are suggestive of subtype 1b to be under less fitness constraints compared to 1a, potentially making it easier for 1b to escape from antibody-mediated immune pressure. Such a difference between the two subtypes is not apparent by comparing simple sequence level statistics such as residue-wise entropies (Supplementary Fig. S7).

### C. Evolutionary modelling and escape time prediction for subtype 1b

To further investigate whether the ease of escaping immune responses is subtype-dependent, we quantified the time it takes for E2 1b to escape immune responses using an in-host viral evolutionary model similar to the one we had employed previously for E2 1a [17]. This evolutionary model takes into account stochastic dynamics during in-host viral evolution, such as host-viral interactions, competition within the viral population, and multiple pathways that may be employed by the virus to escape from immune pressure. This was accomplished by incorporating the inferred E2 1b fitness landscape into a Wright-Fisher-like population genetics model [46]. The parameters involved in this model, such as mutation rate and effective size of the viral population, were set according to known values for HCV [47]–[49], while the sequence survival probability in the population from one generation to the next was determined by the inferred E2 1b fitness landscape (see Methods for details).

For each E2 1b residue, we used the evolutionary model to predict the time for an escape mutation to reach a majority in the population. Specifically, for calculating the escape time associated with a particular residue, we started the evolutionary simulation by first initializing the population with duplicates of a sequence randomly picked from the MSA having the consensus amino acid at the selected residue. Immune pressure was modeled as a fixed reduction in fitness of sequences having the consensus amino acid at the residue, thereby allowing a selective advantage to sequences that incur a mutation at that residue. We continued the simulation until the sequences having a mutation at the selected residue reached a majority in the population. We considered this as a representative marker of viral escape from the corresponding immune pressure. The number of generations for escape was recorded, and this procedure was repeated multiple times with the same initial sequence, as well as with multiple distinct initial sequences. The mean number of generations over all these simulation runs, termed “escape time”, was computed and recorded for every E2 1b residue (see Methods for details).

We validated the predicted escape times for E2 1b residues using experimental and clinical data. First, we assessed the ability of our model to predict known escape mutations from multiple E2-specific HmAbs [39]–[44] (listed in Supplementary Table S1). Such mutations would be expected to be associated with lower escape times compared to mutations at other residues, enabling the virus to escape the associated antibody pressure. Our results demonstrated that this was indeed the case *(P* = 1.4 × 10^-19^, Mann-Whitney test, Fig. 3a). We also validated the escape time metric by predicting the escape times associated with mutating the buried and exposed (surface) residues in E2 1b respectively. Buried residues that form the protein core are more likely to be crucial for stability [50], implying that mutations at these residues would be expected to be associated with higher escape times than mutations at the exposed ones. Residues of the available crystal structure of E2 1b (PDB ID: 6MEI) [51] were classified as exposed or buried according to the standard relative solvent accessibility (RSA) metric (see Methods for details). Comparing the escape times associated with these two sets of residues revealed that the mutations at buried residues were predicted to have higher escape times (*P* = 1.8 × 10^-11^, Mann-Whitney test; Fig. 3b), as expected. Overall, these tests provide confidence in the capability of the evolutionary model to distinguish E2 1b residues associated with low and high escape times.

**Fig. 3:**
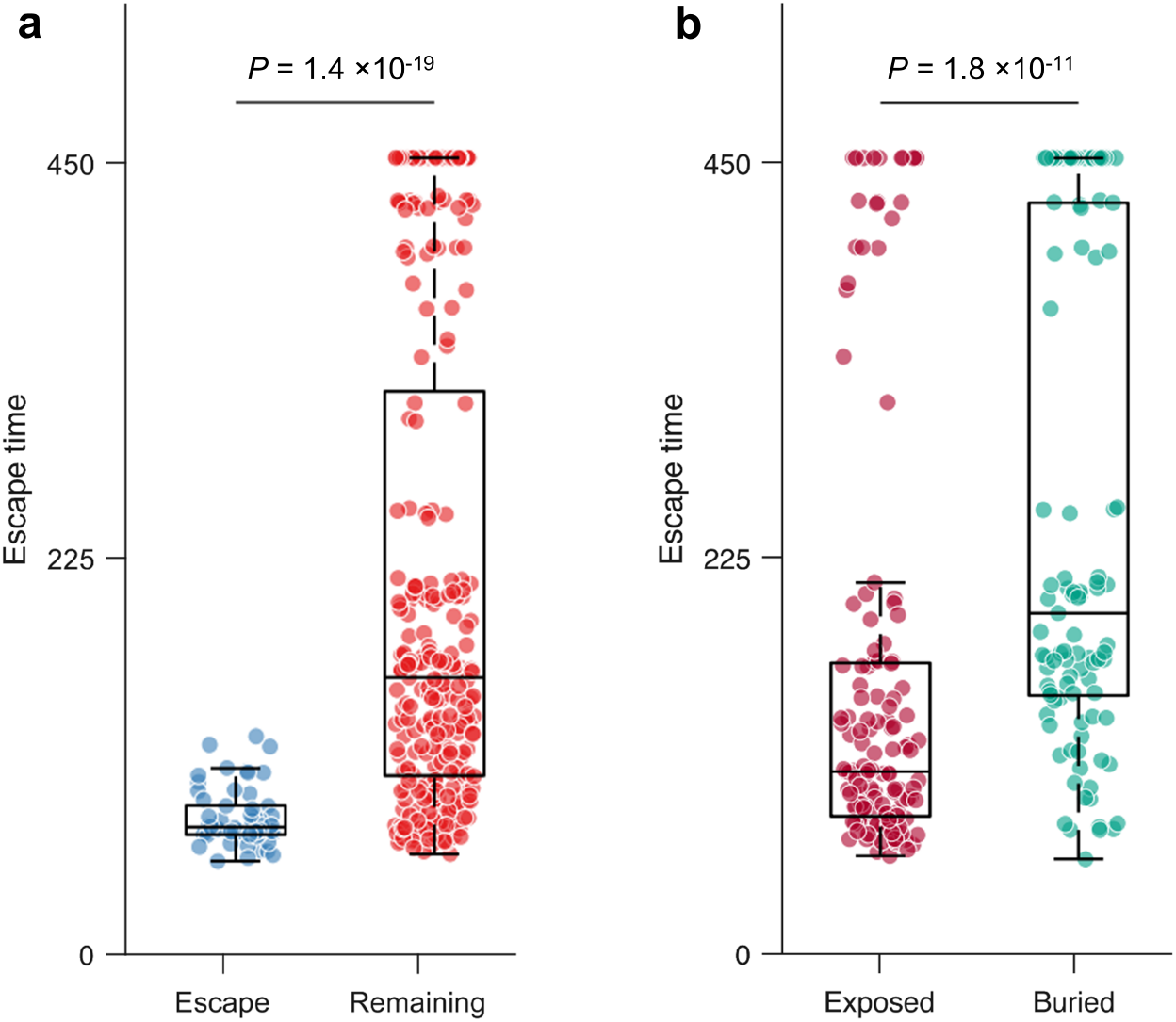
Validation of E2 1b escape times predicted using the in-host evolutionary model. (**a**) Comparison of escape times associated with the residues at which mutation is known to assist in escape from HmAbs (listed in Supplementary Table S1) and those of the remaining E2 residues. (**b**) Comparison of escape times associated with the mutations at exposed and buried residues. Residues of the available crystal structure of E2 1b (PDB ID: 6MEI) [51] were classified as exposed or buried according to the standard RSA metric (see Methods for details). For each box plot, the middle line indicates the median, the edges of the box represent the first and third quartiles, and whiskers extend to span a 1.5 interquartile range from the edges. All reported p-values were calculated using the one-sided Mann-Whitney test.

### D. Escape time comparison indicates subtype 1b may evade HmAbs more easily than subtype 1a

We compared the predicted escape times of the exposed residues of E2 1b (which are more likely to be targeted by HmAbs) with those previously predicted for E2 1a [17]. All relevant model parameters were matched to perform a fair comparison between the two subtypes (see Methods for details). We found that the escape times of exposed residues of E2 1b were generally lower than those of 1a (P = 9.5 × 10^-9^, Mann-Whitney test; Fig. 4a), suggesting that it may be easier for subtype 1b to escape from antibody responses than 1a. This was also true if all residues of both subtypes were considered (P = 1.7 × 10^-13^, Mann-Whitney test; Supplementary Fig. S8). To visualise the escape times associated with exposed residues, we superimposed them as a heat map on the resolved crystal structure of E2 for each subtype (Fig. 4b). Here, as in Fig. 4a, the low concentration of residues with high escape time (coloured in red) in subtype 1b compared to 1a is also evident. Moreover, we observed that few common exposed residues were associated with very high escape times for both subtypes (Supplementary Fig. S9). Interestingly, an epitope 412-425 comprising such residues (423 and 425) was shown in a recent study to be capable of inducing potent neutralizing antibodies against multiple HCV genotypes including 1a and 1b [52]. Hence, targeting these residues may assist in eliciting an effective immune response against both subtypes.

**Fig. 4:**
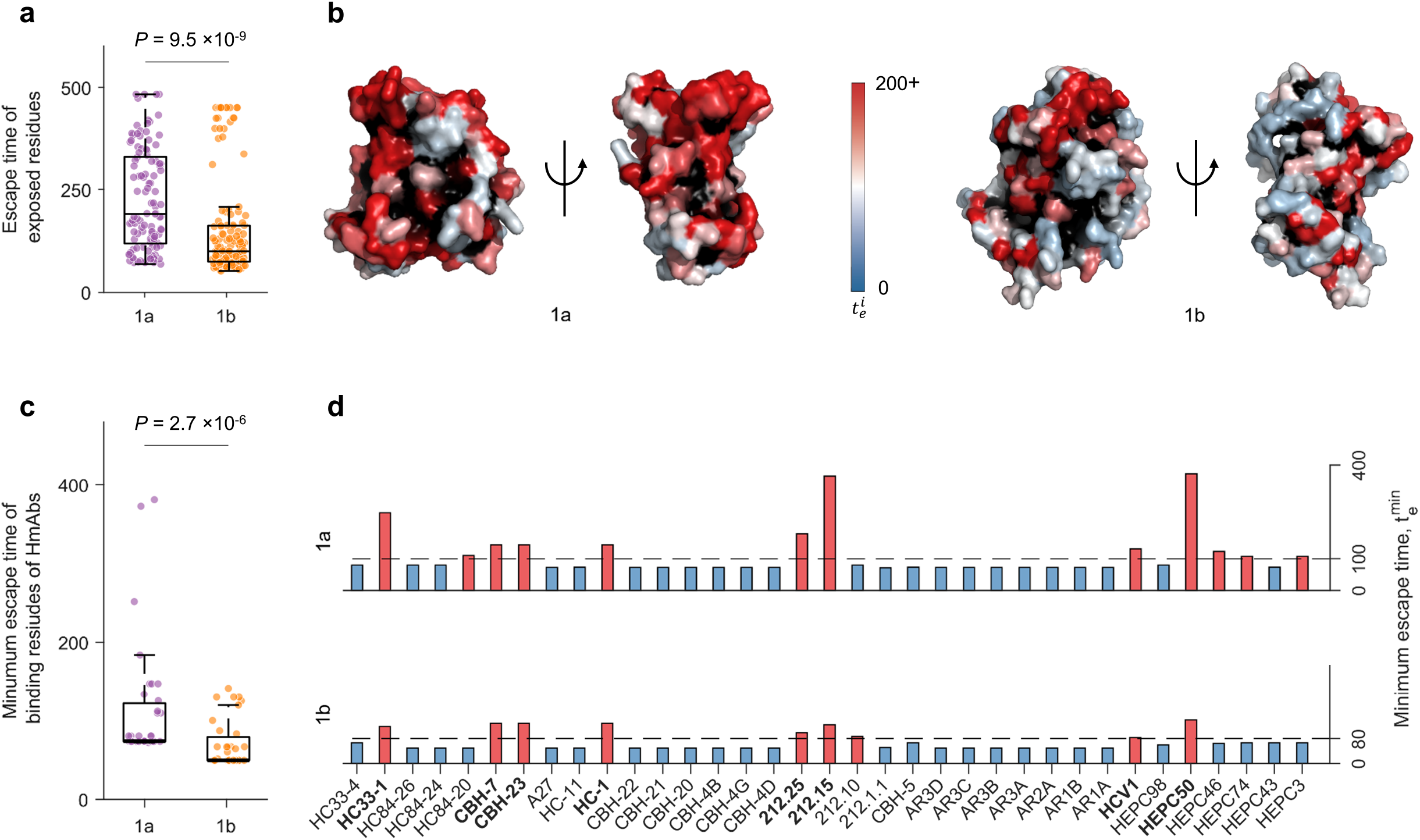
Escape time comparison of E2 subtype 1a and 1b. (**a**) Comparison of escape times associated with exposed residues of E2 of each subtype. (**b**) Escape times of exposed residues superimposed on the crystal structure of E2 1a (left panel, PDB ID: 4MWF [87]) and E2 1b (right panel, PDB ID: 6MEI [51]). Buried residues are shown in black. (**c)-(d**) Comparison of the minimum escape time associated with mutating the binding residues of known E2-specific HmAbs for each subtype (**c**) taken together and (**d**) individually. Binding residues of an antibody were determined by global alanine scanning mutagenesis [53]-[56], where each residue of the wild-type sequence was substituted by alanine (or glycine/serine if the residue in the wildtype was alanine). For the majority of antibodies [53]-[55], the fraction of the mutant sequence binding with respect to the wild-type sequence, called relative binding (RB), was reported for each residue, and we defined binding residues of any one of these antibodies as residues with RB less than or equal to 20%. For some specific antibodies (HEPC98, HEPC50, HEPC46, HEPC74, HEPC43, and HEPC3), critical binding residues as reported in ref. [56] were used. The dashed line for each subtype denotes the optimal cut-off value ζ (see Methods for details). HmAbs predicted to be escape-resistant for any subtype are colored in red, while the remaining ones are colored in blue. The HmAbs predicted to be escape-resistant for both subtypes are shown in bold. In each box plot in (**a**) and (**c**), the middle line indicates the median, the edges of the box represent the first and third quartiles, and whiskers extend to span a 1.5 interquartile range from the edges. All reported p-values were calculated using the one-sided Mann-Whitney test.

We further employed our model to compare the two subtypes based on the escape resistance of known HmAbs. Specifically, we focused on HmAbs with binding residues determined using global alanine scanning experiments [53]–[56]. We compared the minimum escape time (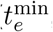, see Methods) predicted by each model for the binding residues of 35 antibodies. Similar to Fig. 4a, this analysis also showed that the minimum escape times predicted for each antibody for subtype 1b were statistically significantly lower than those predicted for 1a (*P* = 2.7 × 10^-6^, Mann-Whitney test; Fig. 4c).

In addition, we considered a linear classifier [17] to compute an optimal escape time cut-off value *ζ* for each subtype to determine whether a residue is relatively escape-resistant or not. This binary classifier was designed by taking the residues with known escape mutations (listed in Supplementary Table S1) as true positives and all remaining residues as true negatives (see Methods for details). We then assessed each antibody by comparing the minimum escape time predicted by each model with the corresponding optimal cut-off value ζ. Based on this analysis, we observed that fewer HmAbs were predicted to be difficult to escape for 1b than 1a (colored in red in Fig. 4d). All the HmAbs predicted to be difficult to escape for 1b were also predicted to be the same for 1a, except for HmAb 212.10. However, the opposite was not true, further suggesting the potentially greater ability for subtype 1b to escape from HmAbs. Nonetheless, we identified eight HmAbs, HC33-1, CBH-7, CBH-23, HC-1, HEPC50, 212.25, 212.15 and HCV1, predicted to be difficult to escape for both subtypes (shown in bold). This result is consistent with information available for these HmAbs in the literature. Specifically, multiple studies have reported HC33-1 as a potentially escape-resistant antibody for multiple genotypes [53], [57] and HCV1 as a potent cross-reactive antibody [58], [59]. Other studies have isolated HmAbs HEPC50, 212.25, and 212.15 from patients who have spontaneously cleared HCV (subtypes 1a, 1b or 3a), and these antibodies have also been shown to be cross-neutralizing [54], [56]. Additionally, among the HmAbs that we predicted to be relatively easy to escape for both subtypes, HmAbs CBH-4D, CBH- 4G, CBH-4B, CBH-20, CBH-21, CBH-22, AR1A, AR1B and AR2A have been reported to be non-neutralizing or isolate-specific antibodies in previous studies [60]–[62]. As for HmAbs AR3A, AR3B, AR3C and AR3D, there are contradicting results in the literature. Certain studies have reported them as potent broadly neutralizing antibodies [62], [63], whereas other studies have found escape mutations against these antibodies [44], [45]. Our results are consistent with the latter set of studies, as both 1a and 1b models predicted these antibodies to be relatively easy to escape.

## III. Discussion

Subtypes 1a and 1b are the two prevalent subtypes of HCV, together representing roughly half of all HCV infections worldwide. Clinical outcomes of subtype 1b infections have been shown to be considerably more severe than those of 1a, showing higher chronicity rate and higher risk of developing cirrhosis and HCC. The underlying reasons for these differences are not well understood. We reasoned that this may be due to subtype 1b being under relatively less fitness constraints than 1a, potentially enabling it to evade immune pressure more easily than 1a. We developed computational models to validate this hypothesis. First, by inferring a fitness landscape of HCV E2 1b and comparing it with an analogous model for 1a obtained previously, our analysis suggested that HCV 1b is subject to less fitness constraints. This result, taken more generally, showed that different subtypes of HCV may possess markedly distinct evolutionary properties. We incorporated the inferred fitness landscape of E2 1b into a population-genetics evolutionary model to predict the time it takes HCV to escape immune responses targeting any specific residue of E2 1b, and compared with the escape times predicted for E2 1a in our previous study [17]. The escape times of E2 1b were generally lower than those of 1a, suggesting that it may be easier for 1b to escape antibody responses.

The significance of incorporating interactions between mutations at different residues has been demonstrated in multiple previous studies for inferring protein fitness landscapes [17], [20], [21], [23]–[25], [37] and for determining networks of residues important for mediating protein structure and function [64]–[68]. However, residue-residue interactions, represented in the maximum entropy fitness landscape model by the coupling parameters (Eq. 3 in Methods), seem to be especially important for HCV E2 1b. This is evident from the significantly higher correspondence between our model predictions and in-vitro fitness (infectivity) measurements 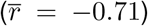, compared with that obtained with a model without couplings 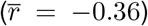 (Fig. 1a). This difference is more apparent than observed previously for models of HIV proteins [21], [22], [24], [25] or other HCV proteins [20] (as well as E2 subtype 1a [17]), suggesting that collective mutational effects, including compensatory mutations, may play a particularly dominant role in shaping the evolution of E2 1b and in providing coordinated evolutionary pathways to facilitate immune escape. Importantly, the incorporation of residue-residue interactions exposed evolutionary differences between the HCV subtypes 1a and 1b (Fig. 2) that were not evident with site-independent models, since at the level of single-site entropy (or single-site genetic variation), both subtypes appeared noticeably similar (Supplementary Fig. S7). Hence, the incorporation of collective mutational effects appears essential to elucidate the distinct evolutionary properties of the different HCV subtypes.

Our results suggest that distinct fitness constraints of HCV subtype 1a and 1b may contribute to the observed differences in disease outcomes of these subtypes. However, other factors may also contribute to disparate disease outcomes, including host-specific factors such as patient’s ethnicity, alcohol consumption, and coinfection with HIV [69]–[71]. Immunogenicity of the viral subtypes may also play a role, since if HCV subtype 1b was found to be less immunogenic than 1a, it could contribute to higher rates of chronic disease. However, there is little evidence to support this, with experimental studies in fact suggesting that the two subtypes have similar immunogenicity [72]. This was demonstrated by stimulating the sera from HCV-infected patients with peptides selected from regions of the HCV proteome having high inter-subtype variability. Similar immunogenicity of both subtypes is also consistent with experimentally-identified HCV E2-specific B cell epitope data available from the Immune Epitope Database (IEDB; https://www.iedb.org) [73], which shows that the coverage of epitopes along the E2 primary structure is similar for both 1a and 1b (Supplementary Fig. S10). Based on current information, it appears unlikely that differences in immunogenicity is a key factor responsible for the disparate disease outcomes of HCV subtypes 1a and 1b.

While to our knowledge possible association between viral fitness constraints and disease outcome of HCV subtypes has not been elucidated previously, related results have been reported for HIV. Like HCV, HIV subtypes have also been shown to associate with different disease outcomes, defined based on the rate of CD4^+^ T cell decline and speed of progression to AIDS [16], [74]–[77]. Clinical studies on HIV have pointed to replicative fitness of the infecting subtype as an important factor associated with disease outcome [16], [78]. For instance, in [16], by following the decay of CD4^+^ T cell counts in untreated patients infected with HIV subtype A, C, or D, patients infected with subtype D were reported to be associated with faster disease progression. Further, by performing dual HIV competition assays of strains isolated from patients infected with each subtype against the reference HIV-1 subtype B isolates, it was shown that the relative replicative fitness of subtype D strains was higher than the other two subtypes. Overall, these studies are supportive of our findings for HCV by providing evidence that associations between the viral subtype and disease progression may be explained, at least in part, by differences in subtype fitness constraints. Our study motivates targeted HCV experimental studies for further investigating the effects of subtype fitness constraints on disease outcome. These may include longitudinal studies of patients infected with HCV subtypes 1a and 1b, which compare the relative fitness (or infectivity) of the infecting strains as well as their association with chronicity or clearance.

The escape times predicted by our in-host evolutionary models identified eight HmAbs (HC33-1, CBH-7, CBH-23, HC-1, HEPC50, 212.25, 212.15 and HCV1) that appear relatively difficult to escape for both subtypes, and also identified specific exposed residues in E2 that are associated with high escape times for both subtypes (Supplementary Fig. S9). Incorporating the epitopes targeted by the identified escape-resistant HmAbs into rationally designed vaccines [79] may aid in eliciting an effective immune response against both HCV subtypes 1a and 1b, and eventually help to curb their global spread. Targeting the exposed yet difficult-to-escape E2 residues, e.g., by synthetic antibodies [80]–[82], may also present an effective therapeutic strategy. Our study motivates the further investigation of the identified escape-resistant antibodies and difficult-to-escape exposed E2 residues in the context of HCV vaccines and therapies via targeted experiments.

## IV. Methods

### A. Data preprocessing

We downloaded 1,559 aligned HCV E2 subtype 1b amino acid sequences (genome coverage ≥ 99%) from the HCV-GLUE database (http://hcv.glue.cvr.ac.uk) [18], [19]. To exclude any outlying sequences, we constructed a pair-wise similarity matrix (1559 × 1559) of the sequences [83], with each (i,j)th entry representing the fraction of residues at which the sequence *i* and sequence *j* are identical. By investigating the first two principal components (PCs) of this matrix, we excluded 260 outlying sequences which appeared at more than 3 scaled median absolute deviations [84] away from the median of either the first or second PC. We also excluded 16 sequences from chimpanzees and 162 sequences having no patient information. This filtering procedure resulted in a total of *M* = 1121 sequences from *W* = 579 patients. In addition, to control residue quality, we excluded 43 residues that were fully conserved. Thus, the final processed MSA comprised M = 1121 sequences and *N* = 320 residues.

### B. Inference of HCV E2 1b fitness landscape

We inferred a maximum entropy model, i.e., the “prevalence landscape”, for E2 1b to serve as a representative of its underlying fitness landscape. The maximum entropy model is a least-biased model that can reproduce the single and double mutant probabilities of the MSA, defined as

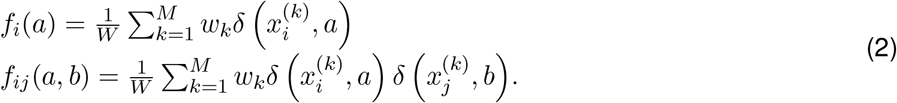

Here, 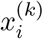 is the *i*th residue of the *k*th sequence from the MSA which takes on a value from either the consensus amino acid 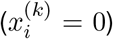 or the mth most frequently observed mutant 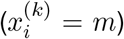 for *m* = 1,…, *q_i_*, where *q_i_* denotes the number of mutants at residue *i. δ* is the Kronecker delta function, *δ*(*a, b*) = 1 if *a* = *b* and 0 otherwise, and *w_k_* is one divided by the number of sequences contributed by the patient from which sequence *k* was obtained. For a given sequence x = [*x*_1_,*x*_2_,…, *x_N_*], the maximum entropy model assigns the probability

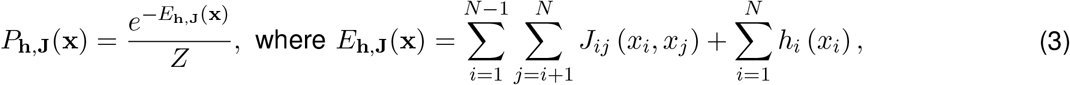

where h is the set of all fields that represent the effect of mutations at a single residue, and J is the set of all couplings that represent the effect of interactions between mutations at two different residues. *Z* = ∑_x_ *e*^-*E*_h,J_(x)^ is a normalization factor, and *E*_h,J_(x) represents the energy of sequence x. The fields h and couplings J are chosen such that the single and double mutant probabilities obtained from the model match the single and double mutant probabilities of the MSA, i.e.,

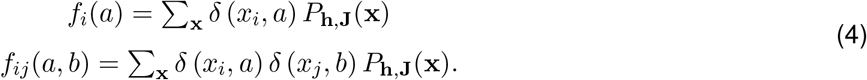

The problem of inferring the model parameters can be cast as the following convex optimization problem [20]

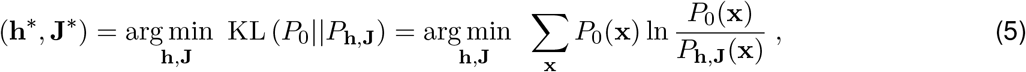

where 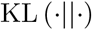 denotes the Kullback-Leibler divergence between probability distributions, and 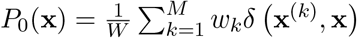 is the patient-weighted probability of observing strain x in the MSA.

As E2 1b is a long protein and has a large residue entropy, the total number of parameters to estimate is very large (Supplementary Fig. S1). To solve this problem, we considered the inference framework, MPF-BML, introduced in [25]. Specifically, we inferred the model parameters using the GUI-based software implementation of MPF-BML [85]. The parameters that we used for model inference are as follows: (i) The sample weights were set according to *w_k_* in Eq. (2); (ii) both *L*_1_ and *L*_2_ regularization parameters were set to 30 for couplings and 10^-3^ for fields, respectively; (iii) the termination condition was set to *ϵ*_1_ <= 2.5 and 0.7 <= ϵ_2_ <= 1.3; and (iv) all other parameters were set to their default values. The statistics of the E2 1b model inferred using these parameters aligned well with those obtained from the MSA (Supplementary Fig. S2).

### C. Fitness verification

We used in-vitro experimental fitness measurements compiled from the literature [27]–[30] to validate that our inferred prevalence landscape is a good proxy of the E2 1b fitness landscape. To normalize for potential experimental biases, we considered the weighted average of Spearman correlation coefficients from different experiments. This is given by

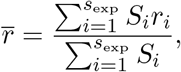

where *S_i_* is the number of measurements and *r_i_* the Spearman correlation coefficient for experiment *i*, and *s*_exp_ is the total number of experiments.

### D. Residue-residue contact prediction

To investigate if the inferred maximum entropy model couplings can predict residue-residue tertiary contacts in the E2 1b protein, we considered the Frobenius norm *F_ij_* of the corresponding coupling matrix *J_ij_* between residue *i* and *j*, a common measure of the strength of residue-residue connections [32], [86]. We then applied an average product correction (APC) to suppress phylogenetic bias and finite sampling effects [33], [86]. These are given by:

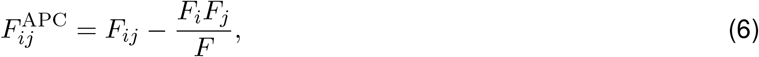

where

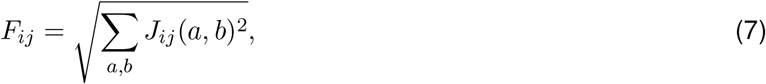

and

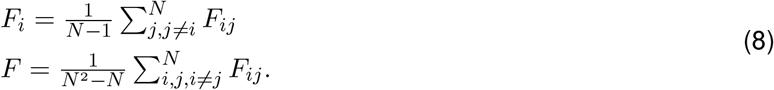

As residues in contact are likely to be strongly coupled, we expected the quantity calculated in Eq. (6) between these residue pairs to be large. The true contacts were determined from the available E2 1b crystal structure (PDB ID: 6MEI) [51], where two residues were assumed to be in contact if their carbon-alpha atoms were less than 8Å apart.

### E. Metrics for comparing the ruggedness of fitness landscapes

We adopted two metrics to compare the fitness landscape of E2 1b with E2 1a [36].

1. *Autocorrelation:* Autocorrelation is used to quantify the average change in fitness as one moves randomly along the landscape. Starting with *M_c_* = 10^6^ random sequences chosen within a Hamming distance *D* = {5,30} from the MSA, we performed a 50-step random walk along the landscape starting from each sequence. The autocorrelation of the sequence energies at the kth step was calculated as

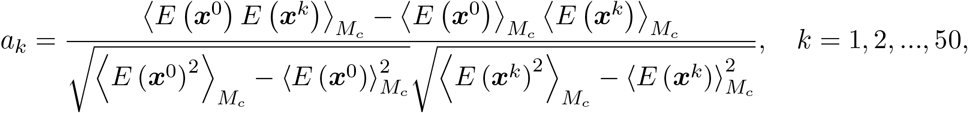

where *x^k^* is the sequence at the kth step, and 〈·〉*_M_c__* is the average over *M_c_* sequences.
2. *Neutrality:* Neutrality quantifies the maximum number of steps (measured in terms of mutations) that one can take in a landscape without much change in fitness, i.e., the resulting difference in fitness remains within a small value e. We started with *M_c_* = 10^6^ random sequences chosen within a Hamming distance *D* = 5 from the MSA. For each starting sequence *x*^0^, we performed an *L* = {500,1000} random walk along the landscape. At each step *k*, we accepted the new sequence *x*^*k*’^ generated by the random walk if the difference in fitness between the new sequence and the current sequence was less than e. Otherwise, we kept the current sequence, i.e., ***x***^*k*+1^ = ***x**^k^*. We calculated the Hamming distance *d^k^* between the sequence ***x**^k^* and the starting sequence ***x***^0^. The neutrality was then calculated as the average over the maximum Hamming distances obtained from all *M_c_* random sequences for a particular *ϵ*, i.e., 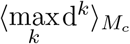.

### F. Comparing the fitness landscapes based on the local peaks

As the local fitness maxima of the landscape, known as local “peaks”, have been shown to be representative of immune escape pathways of HIV [38], we studied the peak structure of the inferred landscapes of subtype 1a and 1b. For each sequence present in the MSA, we obtained the corresponding local peak using the following procedure: For a given sequence of the MSA, we compared the energies of all its neighboring sequences (defined as one Hamming distance away from the sequence), and then chose the most fit sequence (i.e., the sequence with the lowest energy). We repeated the above procedure until the “peak sequence”, i.e., the sequence which has higher fitness than all of its neighbors, was reached. The obtained unique number of local maxima represented the number of local peaks in the specific landscape.

We calculated the statistical significance of enrichment of known escape mutations (listed in Supplementary Table S1) in the peak sequences using a *p*-value. The enrichment of a peak sequence is defined as the fraction of escape mutations among all mutations defining that peak sequence. The *p*-value corresponds to the probability of observing at least *i* mutations out of *j* escape mutations in a peak sequence, where there are *n* total mutations defining the peak sequence out of 363 total residues of the E2 protein. Mathematically, this can be written as

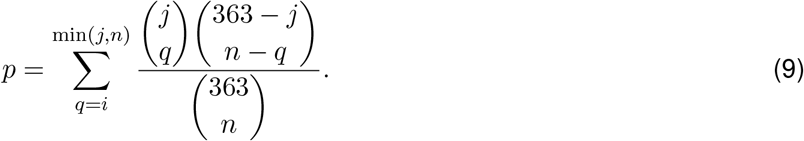

We tested the null hypothesis that these i escape mutations were observed in a peak sequence by a random chance, and it was rejected if *p* < 0.05.

### G. In-host evolutionary model

We considered a population genetics viral evolutionary model similar to [17] (which drew upon an earlier work [23]) for quantifying the ease of escape from antibody responses for each residue in E2 1b. This was accomplished using the “escape time” metric, which represents the number of generations required for mutations at a residue under immune pressure to reach a frequency > 0.5 in a fixed-sized virus population.

Specifically, we employed a well-known population genetics Wright-Fisher model [46]. In this model, sequences in the population undergo mutation, selection, and random sampling steps in each generation. The virus population size was fixed at *M_e_* = 2000, in line with the known HCV effective population size in in-host evolution [47]. For a given E2 1b residue i, we started the simulation with a homogeneous population comprising copies of a sequence randomly selected from the MSA sequences having the consensus amino acid at residue i. In the mutation step, each nucleotide in the sequences was randomly mutated to another nucleotide with a fixed probability *μ* = 10^-4^, in line with the HCV mutation rate reported in [48], [49]. In the selection step, the survival probability of each sequence in the population was calculated based on its fitness predicted from the inferred landscape (see [17] for details). To model the immune pressure at residue *i*, the fitness of all sequences having the consensus amino acid at residue i was decreased by a fixed value *b*, thereby providing a selective advantage to the sequences having a mutation at this residue. Similar to [17], b was set according to the largest value of the field parameter in the inferred landscape. In the sampling step, the new generation of the population was generated through a standard multinomial sampling process parameterized by the survival probabilities calculated in the previous step and *M_e_*. These three steps (mutation, selection and random sampling) were repeated until the number of sequences having a mutation at residue i reached a majority in the population and the number of generations was recorded. This number was considered a representative of the time (generations) taken by the virus to escape the immune pressure at residue *i*. We repeated this procedure multiple times using the same initial sequence, as well as multiple distinct initial sequences. The escape time 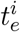 associated with residue *i* was calculated by averaging all number of generations over all these simulation runs.

For a fair comparison of the predicted escape times of E2 1b with those of E2 1a, we used the same parameter values in running the evolutionary simulations for both subtypes. Specifically, we used the same value for the parameter involved in mapping the predicted fitness of a sequence to its survival probability in the population for both 1a and 1b subtypes (see [17]). Moreover, simulations were run for both subtypes for the same number of generations (500), the same number of distinct T/F sequences (25) for each residue, and the same number of simulation runs (100) for each T/F sequence.

### H. Relative solvent accessibility

To determine whether a residue in the crystal structure of each subtype is exposed or buried, we used the get area() function in the PyMOL software (www.pymol.org) with a 1.4 solvent radius parameter to assign each residue in the crystal structure of a subtype (subtype 1a: PDB ID: 4MWF [87]; subtype 1b: PDB ID: 6MEI [51]) with a solvent accessible surface area (SASA). We then obtained the RSA values of each residue by normalizing the respective SASA values per residue in a Gly-X-Gly tripeptide construct [88]. As suggested in [89], residues with RSA > 0.2 were considered as exposed, while the remaining residues were considered buried.

### I. Estimating escape resistance of HmAbs

To compare the escape resistance of known E2-specific HmAbs against each subtype (Fig. 4c, d), we computed the minimum escape time associated with their binding residues (determined using global alanine scanning experiments [53]-[56]). The minimum escape time 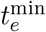 associated with an antibody was defined as

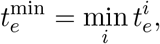

where *i* is selected from the set of binding residues of that antibody. To distinguish whether an E2 residue is relatively escape-resistant or not based on its predicted escape time, we designed a binary classifier using the information of known E2-specific escape mutations (listed in Supplementary Table S1). Specifically, we considered a classifier that takes the residues with known escape mutations as true positives and all remaining residues as true negatives. The classifier for subtype 1b achieved an area under the receiver operating characteristic curve (AUC) of 0.88 (Supplementary Fig. S11a). We chose the optimal cut-off value *ζ* ~ 80 (Supplementary Fig. S11b) based on the maximum F1 score and the maximum Matthews correlation coefficient (MCC)—two commonly used metrics to evaluate the performance of a binary classifier with different thresholds. For subtype 1a, *ζ* was determined to be 100 using a similar statistical analysis in [17]. The HmAbs were classified as relatively escape-resistant for a subtype if their corresponding 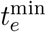 was greater than ζ for that subtype, and vice versa.

## Data and code availability

All data used in this work is publicly available. Accession numbers of E2 1a and 1b sequences used for inferring the models are listed in Supplementary Data 1. The E2 1b infectivity measurements, used for validating the fitness landscape model, are included in Supplementary Data 2. The mean escape time predicted by the in-host evolutionary model for each residue of E2 1b is provided in Supplementary Data 3. The GUI-based software implementation of the MPF-BML method [25], used for inferring the fitness landscape parameters, is available at https://github.com/ahmedaq/MPF-BML-GUI [85]. Data and scripts for reproducing the results are available at https://github.com/hangzhangust/HCVE21a1b_Hang.

## Acknowledgments

The authors were supported by the General Research Fund of the Hong Kong Research Grants Council (RGC) [Grant No. 16204519].

## Supplementary Information

### Supplementary Text

#### S1. Comparison with a conservation-only model

To compare our model with a simpler model based only on amino acid conservation (or single mutant probabilities) that ignores all interactions between residues, we defined a conservation-based maximum entropy model parametrized only by the fields h as

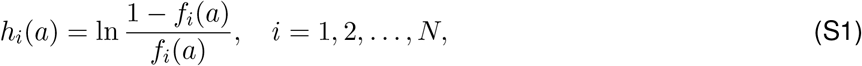

where *f_i_*(*a*) is the frequency of observing amino acid *a* at residue *i*. Correlation between experimental fitness and energy predicted using the conservation-only model was much lower than that of our model (Fig. 1a), suggesting that couplings are significant in determining fitness of HCV strains.

#### S2. Observed local peaks in each landscape are not the result of finite sampling

To test if the local peaks observed in the inferred landscapes are artefacts of finite sampling, we randomly shuffled every column of the MSA of each subtype, which serves to break correlations between residues of the protein while keeping the observed number of amino acids unchanged at each residue. We then inferred a maximum entropy model based on the shuffled MSA of each subtype. Only one local peak was observed in both landscapes, confirming that peaks observed in the inferred fitness landscapes of both E2 1a and 1b were not due to finite sampling effects.

### Supplementary Figures

**Fig. S1:**
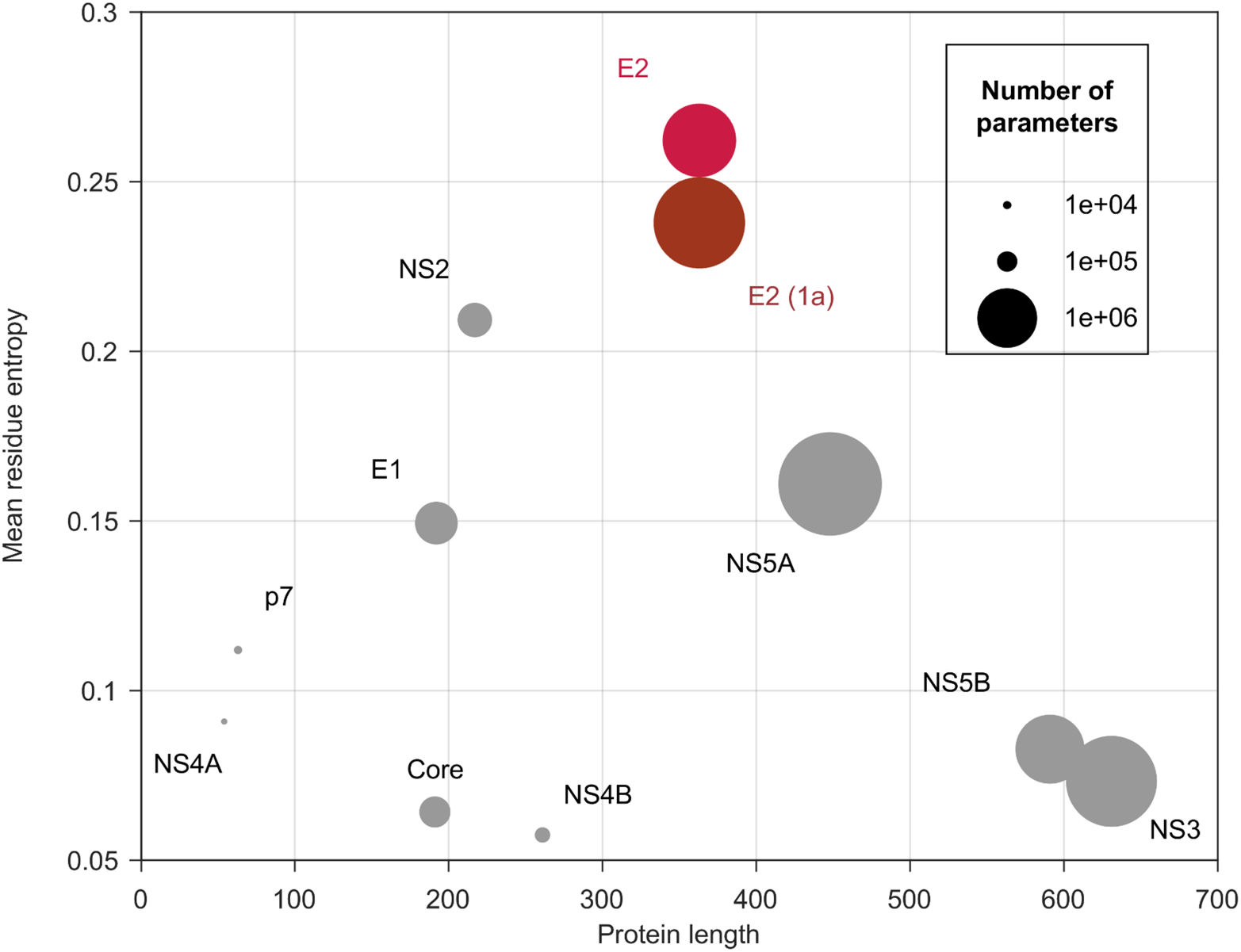
Comparison of the number of parameters required to estimate a fitness landscape for different HCV proteins of subtype 1b. E2 is a long protein with the highest mean residue entropy among HCV proteins. Thus inferring its landscape requires estimating a large number of model parameters. Mean residue entropy *H* can be calculated as 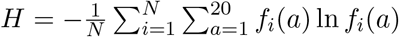, where *f_i_*(*a*) is the frequency of observing amino acid a at residue *i*, and *N* is the number of residues in that protein. The number of parameters was calculated by considering all amino acid mutants observed at each residue. The corresponding numbers for E2 subtype 1a have also been included for reference.

**Fig. S2:**
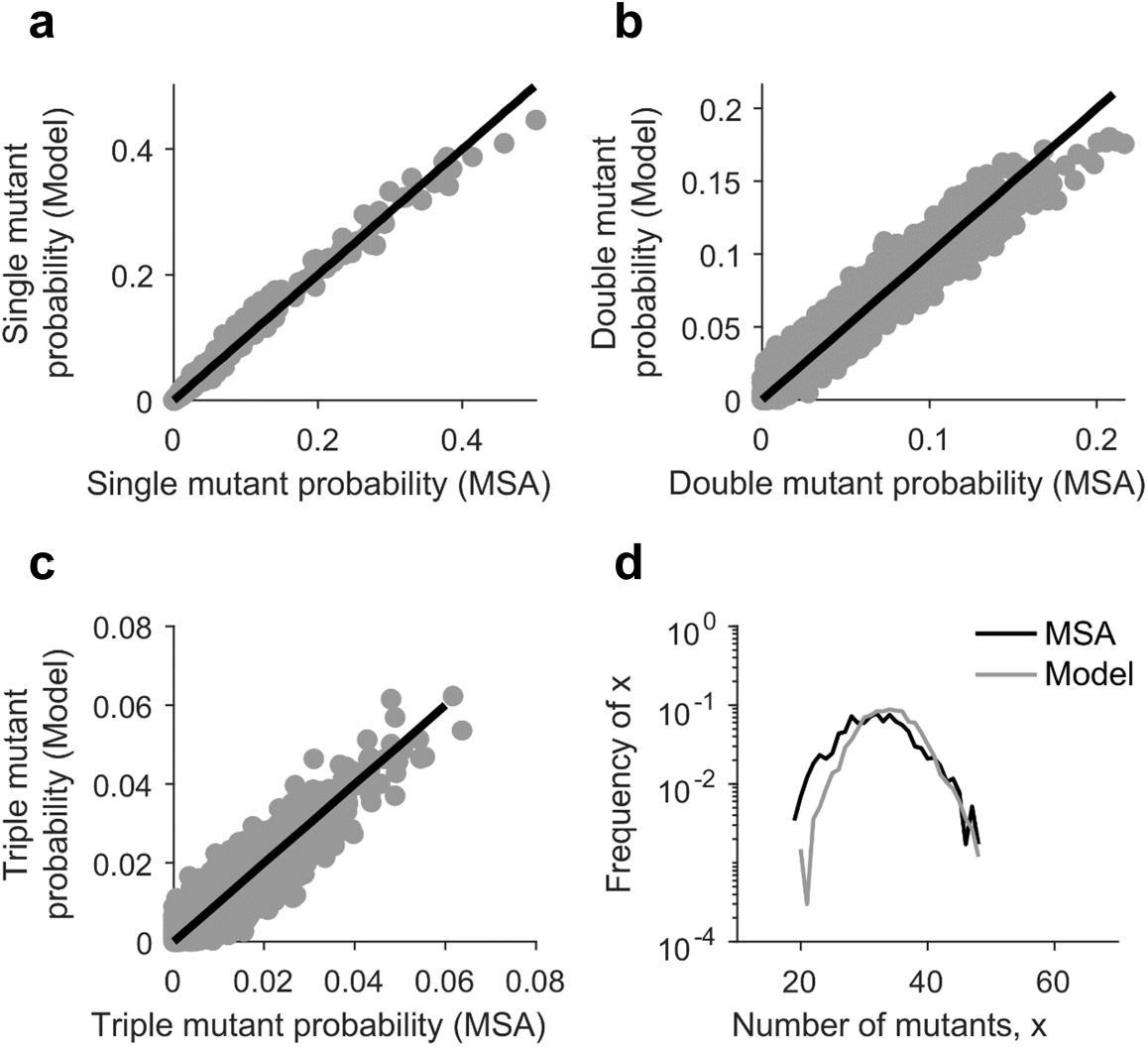
Statistical validation of the inferred E2 subtype 1b landscape. Comparison of the (**a**) single mutant probabilities, (**b**) double mutant probabilities, (**c**) triple mutant probabilities, and (**d**) distribution of the number of mutants per sequence obtained from the MSA and those predicted by the inferred model. Samples were generated from the inferred model using a Markov Chain Monte Carlo (MCMC) procedure [1].

**Fig. S3:**
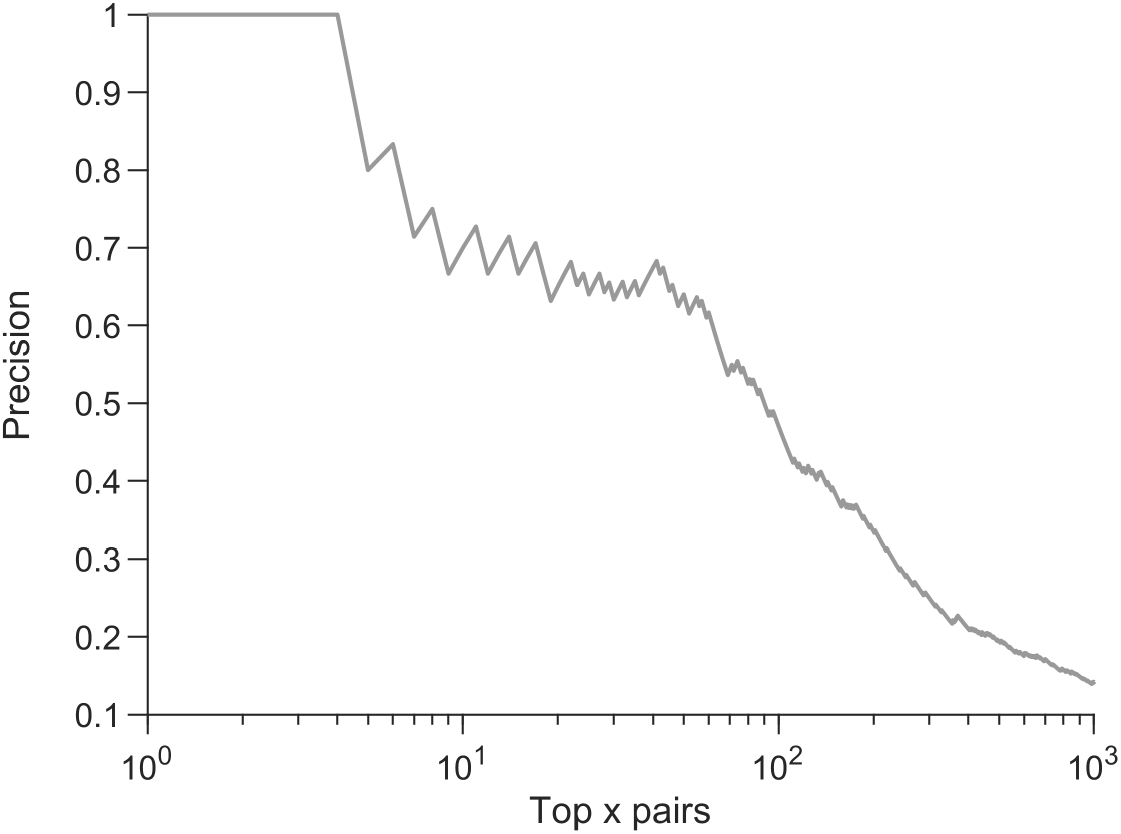
Precision of contact predictions vs. the top x pairs obtained using DCA [2]. Predictions of DCA were generated using the code provided in ref. [2], where the re-weighting procedure was implemented according to Eq. (2) and the pseudocount parameter was set to 0.5. Precision is the proportion of top x pairs that are in contact in the protein tertiary structure. Two residues were assumed to be in contact if their carbon-alpha atoms were <8Å apart according to the available crystal structure of E2 1b (PDB ID: 6MEI).

**Fig. S4:**
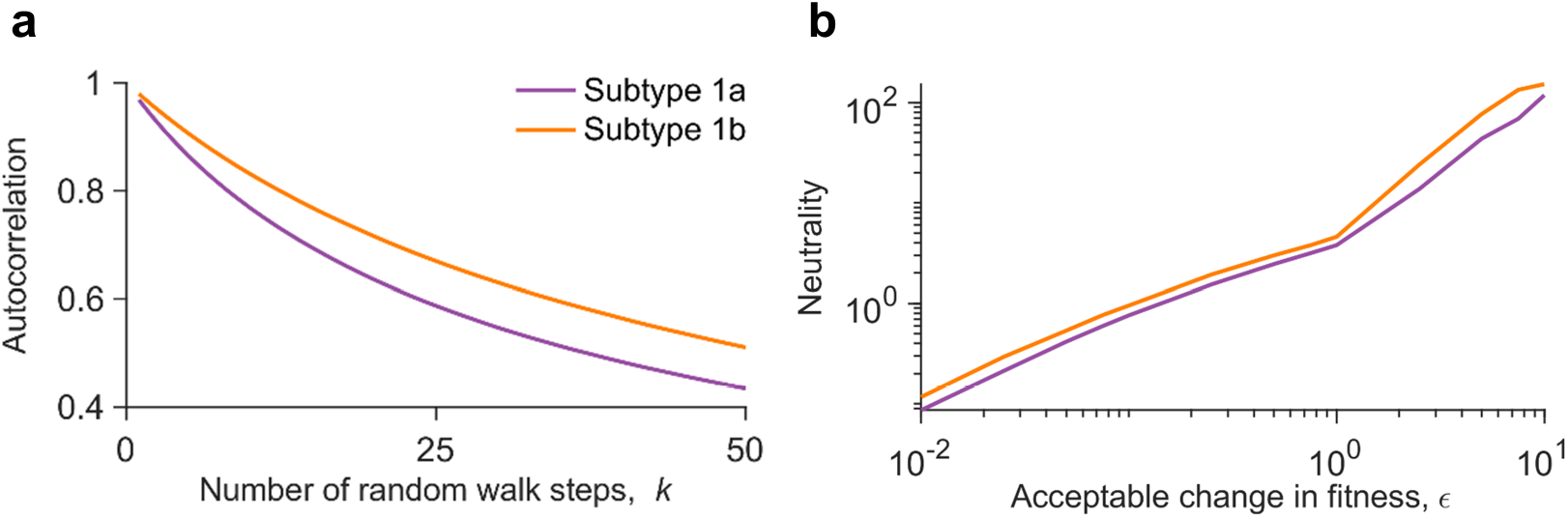
Robustness of the autocorrelation and neutrality results (Fig. 2a) to variation of the involved parameters. (**a**) Comparison of the autocorrelation of sequence energies of the E2 1a and 1b landscapes, with the starting sequences chosen within a Hamming distance (*D*) of 30 from the MSA (see Methods for details). (**b**) Comparison of the neutrality of the E2 1a and 1b landscapes. Neutrality was computed for L = 1000 steps (see Methods for details). It is observed that the fitness constraints on E2 1b appear lower than E2 1a, independently of whether we use *D* = 5 (Fig. 2a, left panel) or *D* = 30 for computing landscape autocorrelation, and whether we use *L* = 500 (Fig. 2a, right panel) or *L* = 1000 for computing landscape neutrality.

**Fig. S5:**
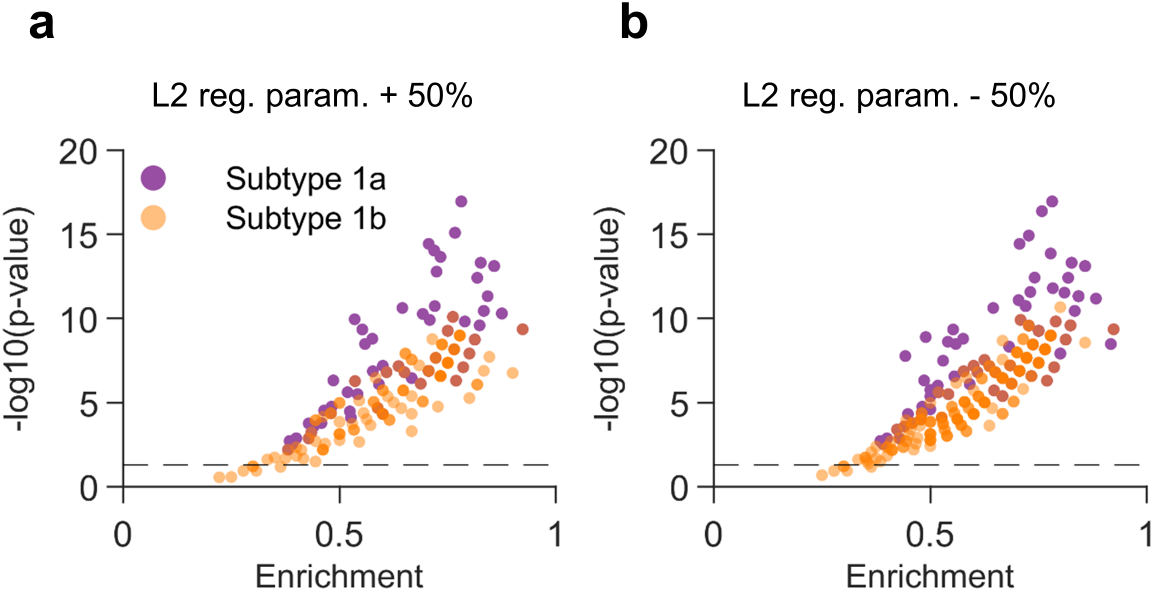
Robustness of the statistical enrichment of known escape mutations among landscape peak sequences (Fig. 2b left panel) with respect to changes in landscape inference parameters. Almost all peak sequences of each subtype are statistically significantly enriched in escape mutations whether the *L*_2_ regularization parameters used in the landscape inference were (**a**) increased by 50% or (**b**) decreased by 50%.

**Fig. S6:**
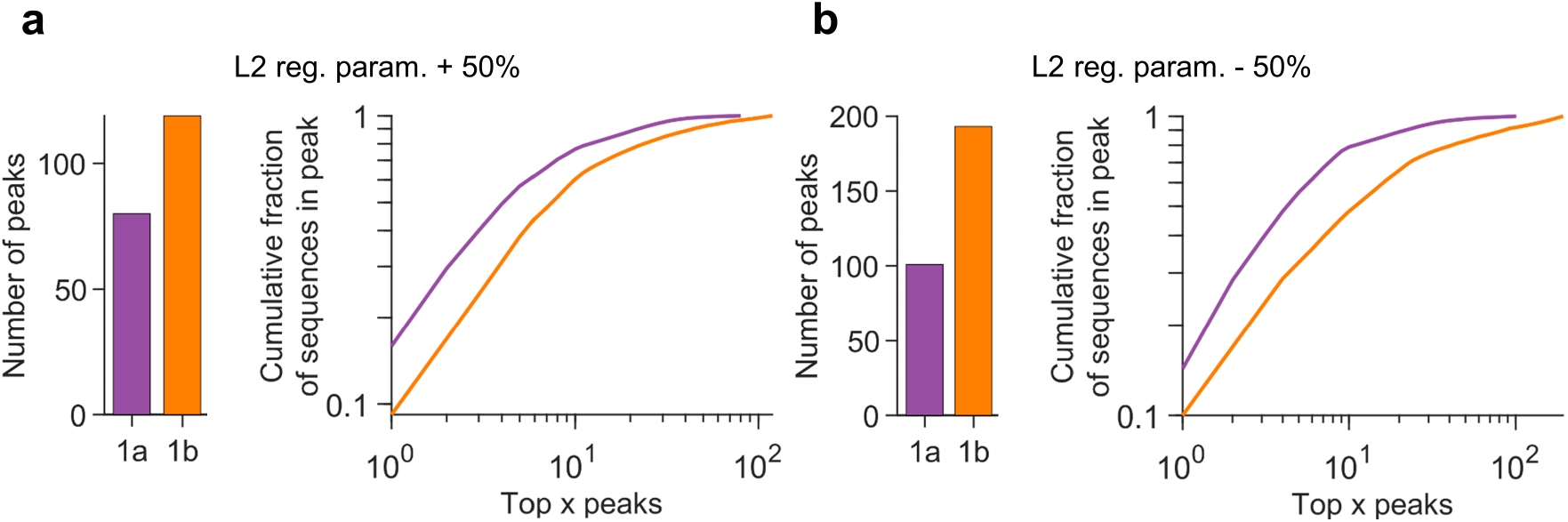
Robustness of the statistics of the observed peaks (Fig. 2b middle and right panels) with respect to changes in landscape inference parameters. (Left panel) Number of peaks and (right panel) cumulative fraction of sequences in peaks observed in each landscape. More number of peaks are observed in 1b landscape than 1a, and peaks are more spread out across peaks in 1b than 1a whether the *L*_2_ regularization parameters used in the landscape inference were (**a**) increased by 50% or (**b**) decreased by 50%.

**Fig. S7:**
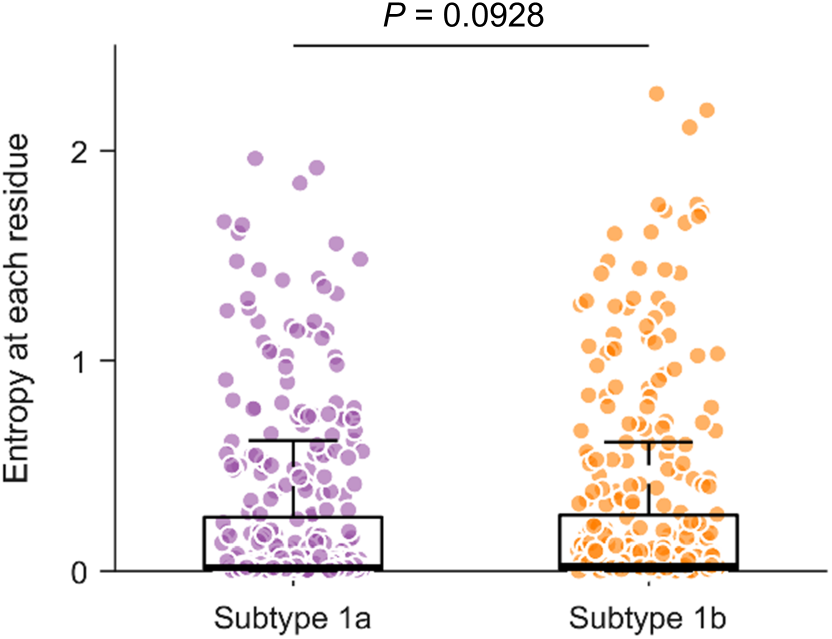
Comparison of the residue-wise entropy of E2 subtypes 1a and 1b. The residue-wise entropy of E2 1a and 1b is not statistically significant different (*P* > 0.05). In each box plot, the middle line indicates the median, the edges of the box represent the first and third quartiles, and whiskers extend to span a 1.5 interquartile range from the edges. The reported *p*-value was calculated using the one-sided Mann-Whitney test.

**Fig. S8:**
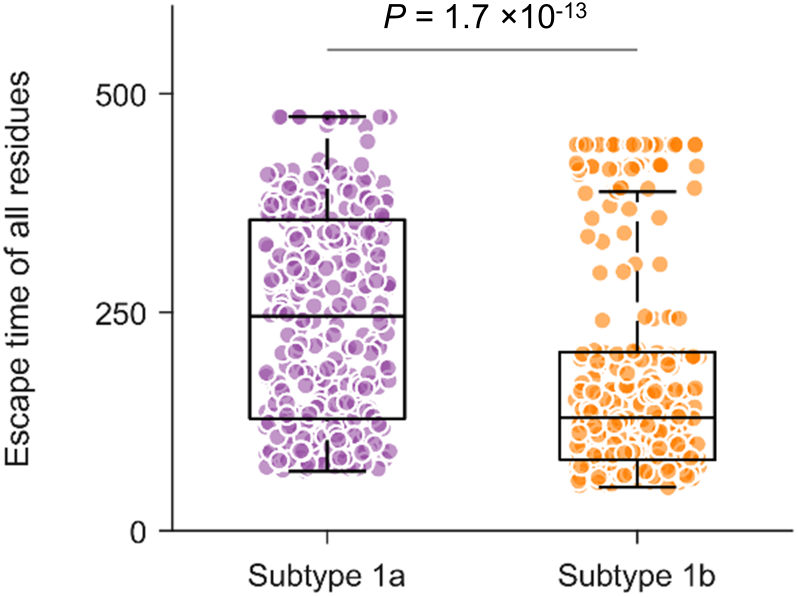
Comparison of predicted escape times of all residues of E2 1a and 1b. Escape times associated with E2 1b are statistically significantly lower than those of E2 1a. In each box plot, the middle line indicates the median, the edges of the box represent the first and third quartiles, and whiskers extend to span a 1.5 interquartile range from the edges. The reported p-value was calculated using the one-sided Mann-Whitney test.

**Fig. S9:**
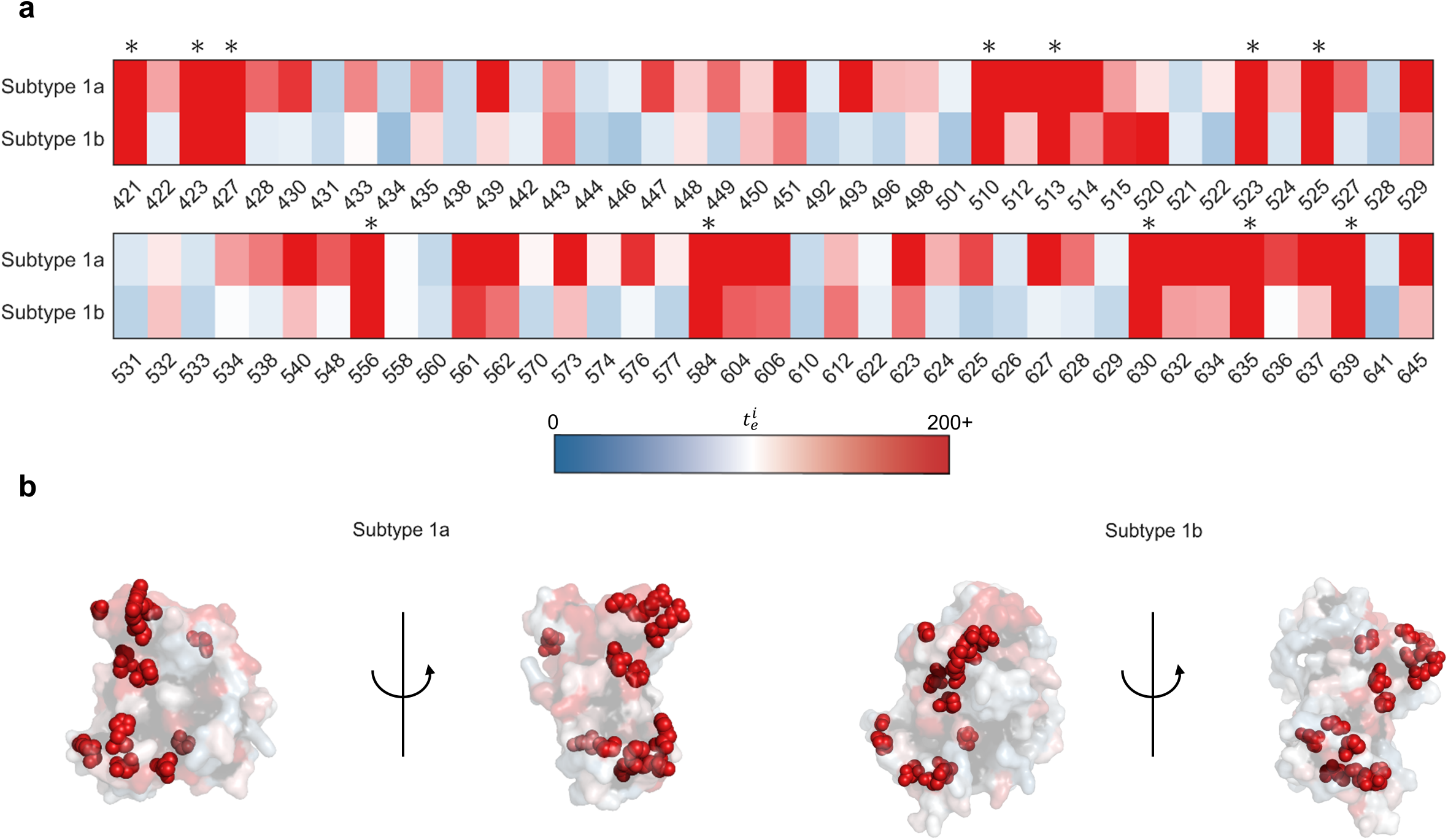
Exposed residues identified to be associated with high escape times for both subtypes 1a and 1b. (**a**) Escape times associated with the E2 residues exposed in both subtypes, with those residues associated high escape times for both subtypes marked with an asterisk on the top of each panel. The x-axis denotes the residue number (according to the H77 reference sequence). While the optimal cut-off value *ζ* (see Methods for details) to determine whether a residue is difficult to escape or not was 100 generations for 1a and 80 generations for 1b, we selected a conservative minimum escape time of 200 to determine exposed residues associated with high escape time for both subtypes. (**b**) Common exposed residues predicted to be associated with high escape time for both subtypes (in (**a**)) are shown as spheres on the crystal structure of E2 1a (left panel, PDB ID: 4MWF) and E2 1b (right panel, PDB ID: 6MEI).

**Fig. S10:**
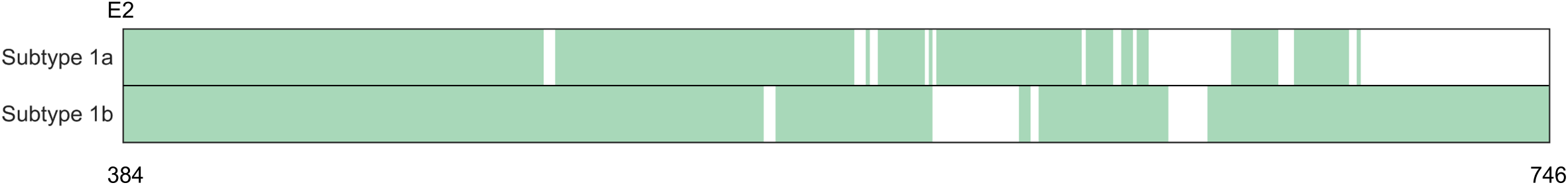
B cell epitope coverage of E2 for subtype 1a and 1b. The experimentally-identified B cell epitopes cover a similar fraction of E2 along its primary structure for both subtypes. The epitopes for each subtype were obtained from IEDB (https://www.iedb.org) [3]. Epitopes corresponding to positive B cell assays of HCV were searched, and E2-specific epitopes for which subtype 1a or 1b had been specified were then filtered. This procedure provided 71 unique B cell epitopes for E2 1a and 33 unique ones for 1b. E2 1a epitopes were reported by 26 independent studies while those for 1b were reported by 7 studies, indicating that the high number for E2 1a epitopes was possibly due to this subtype being studied more in the literature. E2 residues covered by the B cell epitopes are shown in green while the remaining ones are shown in white. Residues are numbered according to the H77 reference sequence.

**Fig. S11:**
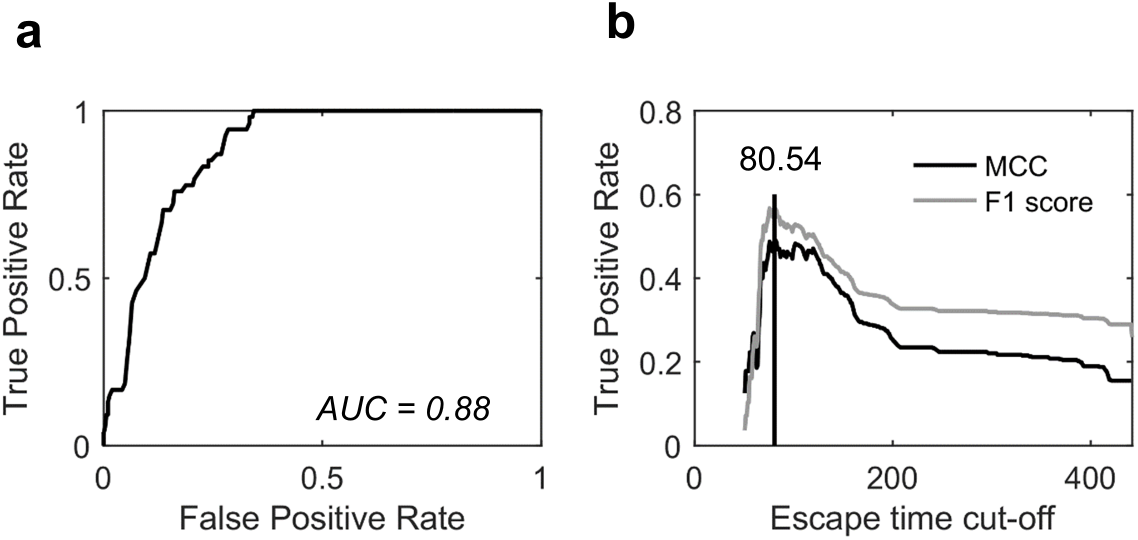
Classifier for determining optimal escape time for E2 1b, designed based on knowledge of experimentally or clinically identified escape mutations. (**a**) Receiver operating characteristic curve for identifying the known escape mutations (listed in Supplementary Table S1) using the escape time metric. (**b**) Determination of the optimal escape time cut-off based on the F1 score and MCC. In this classification, residues with known escape mutations (listed in Supplementary Table S1) were considered as true positives while all remaining residues were assumed true negatives.

### Supplementary Table

**TABLE S1:** List of known escape mutations from E2-specific HmAbs.

| Escape residues | HmAbs | Reference |
| --- | --- | --- |
| 408 | HC33-4 | [4] |
| 384, 386, 388, 390, 391, 393, 394, 395, 396, 397, 398, 399, 400, 401, 402, 403, 404, 405, 407, 410 | HmAbs targeting HVR1 | [5] |
| 431 | CBH-2 | [6] |
| 431, 435, 444, 446, 466, 482, 501, 528, 531, 538, 580, 610, 636, 713 | CBH-8C, CBH-2, CBH-5, HC-2, HC-11 | [7] |
| 391, 394, 401, 415, 417, 434, 444, 608 | HCV1 | [8] |
| 416, 422, 424, 431, 433, 438, 442, 446, 453, 456, 461, 475, 482, 520*, 524, 531, 533, 557, 558, 560 | CBH-2, CBH-5, HC84-22, HC84-26, AR3A, AR3B, AR3C, AR3D | [9] |
| 431, 438, 442 | AR3A | [10] |

\*Excluded from our analysis, as no mutation was observed at this residue in the E2 1b sequence data.

#### Description of Additional Supplementary Files

File Name: Supplementary Data 1

Description: Accession numbers of E2 1a and 1b sequences used for inferring the models.

File Name: Supplementary Data 2

Description: The E2 1b infectivity measurements, used for validating the fitness landscape model.

File Name: Supplementary Data 3

Description: The mean escape time predicted by the in-host evolutionary model for each residue of E2 1b.

## References

[1] H. R. Rosen, “Clinical practice. Chronic hepatitis C infection.” The New England Journal of Medicine, vol. 364, no. 25, pp. 2429–2438, 2011.

[2] A. Petruzziello, S. Marigliano et al., “Global epidemiology of hepatitis C virus infection: An up-date of the distribution and circulation of hepatitis C virus genotypes,” World Journal of Gastroenterology, vol. 22, no. 34, p. 7824, 2016.

[3] Centers for Disease Control and Prevention, “Hepatitis C questions and answers for the public,” 2018. https://www.cdc.gov/hepatitis/HCV/cfaq.htm

[4] E. S. Rosenthal and C. S. Graham, “Price and affordability of direct-acting antiviral regimens for hepatitis C virus in the united states,” Infectious Agents and Cancer, vol. 11, no. 1, p. 24, 2016.

[5] World Health Organization, “Hepatitis C, Fact sheet,” 2019. https://www.who.int/news-room/fact-sheets/detail/hepatitis-c

[6] C. Rossi, Z. A. Butt et al., “Hepatitis C virus reinfection after successful treatment with direct-acting antiviral therapy in a large population-based cohort,” Hepatology, vol. 69, no. 5, pp. 1007–1014, 2018.

[7] D. L. Wyles and A. F. Luetkemeyer, “Understanding hepatitis C virus drug resistance: Clinical implications for current and future regimens,” Topics in Antiviral Medicine, vol. 25, no. 3, pp. 103–109, 2017.

[8] D. B. Smith, J. Bukh et al., “Expanded classification of hepatitis C virus into 7 genotypes and 67 subtypes: Updated criteria and genotype assignment web resource,” Hepatology, vol. 59, no. 1, pp. 318–327, 2013.

[9] J. P. Messina, I. Humphreys et al., “Global distribution and prevalence of hepatitis C virus genotypes,” Hepatology, vol. 61, no. 1, pp. 77–87, 2014.

[10] P. Amoroso, M. Rapicetta et al., “Correlation between virus genotype and chronicity rate in acute hepatitis C,” Hepatology, vol. 28, no. 6, pp. 939–944, 1998.

[11] A. R. Osella, G. Misciagna et al., “Hepatitis C virus genotypes and risk of cirrhosis in southern Italy,” Clinical Infectious Diseases, vol. 33, no. 1, pp. 70–75, 2001.

[12] E. Silini, R. Bottelli et al., “Hepatitis C virus genotypes and risk of hepatocellular carcinoma in cirrhosis: A case-control study,” Gastroenterology, vol. 111, no. 1, pp. 199–205, 1996.

[13] S. Bruno, A. Crosignani et al., “Hepatitis C virus genotype 1b as a major risk factor associated with hepatocellular carcinoma in patients with cirrhosis: A seventeen-year prospective cohort study,” Hepatology, vol. 46, no. 5, pp. 1350–1356, 2007.

[14] S. Raimondi, S. Bruno et al., “Hepatitis C virus genotype 1b as a risk factor for hepatocellular carcinoma development: A metaanalysis,” Hepatology, vol. 50, no. 6, pp. 1142–1154, 2009.

[15] M.-H. Lee, H.-I. Yang et al., “Hepatitis C virus genotype 1b increases cumulative lifetime risk of hepatocellular carcinoma,” International Journal of Cancer, vol. 135, no. 5, pp. 1119–1126, 2014.

[16] C. M. Venner, I. Nankya et al., “Infecting HIV-1 subtype predicts disease progression in women of sub-saharan africa,” EBioMedicine, vol. 13, pp. 305–314, 2016.

[17] A. A. Quadeer, R. H. Y Louie, and M. R. Mckay, “Identifying immunologically-vulnerable regions of the HCV E2 glycoprotein and broadly neutralizing antibodies that target them,” Nature Communications, vol. 10, no. 1, p. 2073, 2019.

[18] J. B. Singer, E. C. Thomson et al., “GLUE: A flexible software system for virus sequence data,” BMC Bioinformatics, vol. 19, no. 1, p. 532, 2018.

[19] J. Singer, E. Thomson et al., “Interpreting viral deep sequencing data with GLUE,” Viruses, vol. 11, no. 4, p. 323, 2019.

[20] G. R. Hart and A. L. Ferguson, “Empirical fitness models for hepatitis C virus immunogen design,” Physical Biology, vol. 12, no. 6, p. 066006, 2015.

[21] A. L. Ferguson, J. K. Mann et al., “Translating HIV sequences into quantitative fitness landscapes predicts viral vulnerabilities for rational immunogen design,” Immunity, vol. 38, no. 3, pp. 606–617, 2013.

[22] J. K. Mann, J. P. Barton et al., “The fitness landscape of HIV-1 Gag: Advanced modeling approaches and validation of model predictions by in vitro testing,” PLoS Computational Biology, vol. 10, no. 8, p. e1003776, 2014.

[23] J. P. Barton, N. Goonetilleke et al., “Relative rate and location of intra-host HIV evolution to evade cellular immunity are predictable,” Nature Communications, vol. 7, p. 11660, 2016.

[24] W. F. Flynn, A. Haldane et al., “Inference of epistatic effects leading to entrenchment and drug resistance in HIV-1 protease,” Molecular Biology and Evolution, vol. 34, no. 6, pp. 1291–1306, 2017.

[25] R. H. Y. Louie, K. J. Kaczorowski et al., “Fitness landscape of the human immunodeficiency virus envelope protein that is targeted by antibodies,” Proceedings of the National Academy of Sciences, vol. 115, no. 4, pp. E564–E573, 2018.

[26] D. K. Murakowski, J. P. Barton et al., “Adenovirus-vectored vaccine containing multidimensionally conserved parts of the HIV proteome is immunogenic in rhesus macaques,” Proceedings of the National Academy of Sciences, vol. 118, no. 5, p. e2022496118, 2021.

[27] L. Esteban-Riesco, F. Depaulis et al., “Rapid and sustained autologous neutralizing response leading to early spontaneous recovery after HCV infection,” Virology, vol. 444, no. 1–2, pp. 90–99, 2013.

[28] R. A. Urbanowicz, C. P. Mcclure et al., “A diverse panel of hepatitis C virus glycoproteins for use in vaccine research reveals extremes of monoclonal antibody neutralization resistance,” Virology, vol. 90, no. 7, pp. 3288–3301, 2015.

[29] A. S. Naik, A. Owsianka et al., “Reverse epitope mapping of the E2 glycoprotein in antibody associated hepatitis C virus,” PLoS ONE, vol. 12, no. 5, pp. 1–20, 2017.

[30] H. Pantua, J. Diao et al., “Glycan shifting on hepatitis C virus (HCV) E2 glycoprotein is a mechanism for escape from broadly neutralizing antibodies,” Journal of Molecular Biology, vol. 425, no. 11, pp. 1899–1914, 2013.

[31] M. Parera and M. A. Martinez, “Strong epistatic interactions within a single protein,” Molecular Biology and Evolution, vol. 31, no. 6, pp. 1546–1553, 2014.

[32] M. Weigt, R. A. White et al., “Identification of direct residue contacts in protein-protein interaction by message passing,” Proceedings of the National Academy of Sciences, vol. 106, no. 1, pp. 67–72, 2008.

[33] S. Dunn, L. Wahl, and G. Gloor, “Mutual information without the influence of phylogeny or entropy dramatically improves residue contact prediction,” Bioinformatics, vol. 24, no. 3, pp. 333–340, 2007.

[34] F. Morcos, A. Pagnani et al., “Direct-coupling analysis of residue coevolution captures native contacts across many protein families,” Proceedings of the National Academy of Sciences, vol. 108, no. 49, pp. E1293–E1301, 2011.

[35] V. K. Vassilev, T. C. Fogarty, and J. F Miller, “Smoothness, ruggedness and neutrality of fitness landscapes: From theory to application,” Natural Computing Series Advances in Evolutionary Computing, p. 344, 2003.

[36] R. D. Kouyos, G. E. Leventhal et al., “Exploring the complexity of the HIV-1 fitness landscape,” PLoS Genetics, vol. 8, no. 3, p. e1002551, 2012.

[37] A. A. Quadeer, J. P. Barton et al., “Deconvolving mutational patterns of poliovirus outbreaks reveals its intrinsic fitness landscape,” Nature Communications, vol. 11, no. 1, p. 377, 2020.

[38] J. P. Barton, M. Kardar, and A. K. Chakraborty, “Scaling laws describe memories of host-pathogen riposte in the HIV population,” Proceedings of the National Academy of Sciences, vol. 112, no. 7, pp. 1965–1970, 2015.

[39] Z.-Y Keck, C. Girard-Blanc et al., “Antibody response to hypervariable region 1 interferes with broadly neutralizing antibodies to hepatitis C virus,” Virology, vol. 90, no. 6, pp. 3112–3122, 2016.

[40] N. Kato, H. Sekiya et al., “Humoral immune response to hypervariable region 1 of the putative envelope glycoprotein (gp70) of hepatitis C virus.” Virology, vol. 67, no. 7, pp. 3923–3930, 1993.

[41] Z.-Y Keck, O. Olson et al., “A point mutation leading to hepatitis C virus escape from neutralization by a monoclonal antibody to a conserved conformational epitope,” Virology, vol. 82, no. 12, pp. 6067–6072, 2008.

[42] Z.-Y Keck, S. H. Li et al., “Mutations in hepatitis C virus E2 located outside the CD81 binding sites lead to escape from broadly neutralizing antibodies but compromise virus infectivity,” Virology, vol. 83, no. 12, pp. 6149–6160, 2009.

[43] T. J. Morin, T. J. Broering et al., “Human monoclonal antibody HCV1 effectively prevents and treats HCV infection in chimpanzees,” PLoS Pathogens, vol. 8, no. 8, p. e1002895, 2012.

[44] J. R. Bailey, L. N. Wasilewski et al., “Naturally selected hepatitis C virus polymorphisms confer broad neutralizing antibody resistance,” Journal of Clinical Investigation, vol. 125, no. 1, pp. 437–447, 2015.

[45] R. Velázquez-Moctezuma, A. Galli et al., “Hepatitis C virus escape studies of human antibody AR3A reveal a high barrier to resistance and novel insights on viral antibody evasion mechanisms,” Virology, vol. 93, no. 4, pp. e01 909–18, 2019.

[46] W. J. Ewens, “Mathematical population genetics,” Interdisciplinary Applied Mathematics, 2004.

[47] R. A. Bull, F Luciani et al., “Sequential bottlenecks drive viral evolution in early acute hepatitis C virus infection,” PLoS Pathogens, vol. 7, no. 9, pp. 1–14, 2011.

[48] J. M. Cuevas, F. Gonzalez-Candelas et al., “Effect of ribavirin on the mutation rate and spectrum of hepatitis C virus in vivo,” Virology, vol. 83, no. 11, pp. 5760–5764, 2009.

[49] R. Sanjuan, M. R. Nebot et al., “Viral mutation rates,” Virology, vol. 84, no. 19, pp. 9733–9748, 2010.

[50] H. Chen, “Prediction of solvent accessibility and sites of deleterious mutations from protein sequence,” Nucleic Acids Research, vol. 33, no. 10, pp. 3193–3199, 2005.

[51] A. I. Flyak, S. Ruiz et al., “HCV broadly neutralizing antibodies use a CDRH3 disulfide motif to recognize an E2 glycoprotein site that can be targeted for vaccine design,” Cell Host & Microbe, vol. 24, no. 5, pp. 703–716.e3, 2018.

[52] A. Czarnota, A. Offersgaard et al., “Specific antibodies induced by immunization with hepatitis B virus-like particles carrying hepatitis C virus envelope glycoprotein 2 epitopes show differential neutralization efficiency,” Vaccines, vol. 8, no. 2, pp. 1–19, 2020.

[53] B. G. Pierce, Z.-Y. Keck et al., “Global mapping of antibody recognition of the hepatitis C virus E2 glycoprotein: Implications for vaccine design,” Proceedings of the National Academy of Sciences, vol. 113, no. 45, pp. E6946–E6954, 2016.

[54] Z.-Y. Keck, B. G. Pierce et al., “Broadly neutralizing antibodies from an individual that naturally cleared multiple hepatitis C virus infections uncover molecular determinants for E2 targeting and vaccine design,” PLoS Pathogens, vol. 15, no. 5, p. e1007772, 2019.

[55] R. Gopal, K. Jackson et al., “Probing the antigenicity of hepatitis C virus envelope glycoprotein complex by high-throughput mutagenesis,” PLoS Pathogens, vol. 13, no. 12, p. e1006735, 2017.

[56] J. R. Bailey, A. I. Flyak et al., “Broadly neutralizing antibodies with few somatic mutations and hepatitis C virus clearance,” JCI Insight, vol. 2, no. 9, p. e92872, 2017.

[57] Z.-Y. Keck, A. G. N. Angus et al., “Non-random escape pathways from a broadly neutralizing human monoclonal antibody map to a highly conserved region on the hepatitis C virus E2 glycoprotein encompassing amino acids 412-423,” PLoS Pathogens, vol. 10, no. 8, pp. 1–13, 2014.

[58] L. Kong, E. Giang et al., “Structural basis of hepatitis C virus neutralization by broadly neutralizing antibody HCV1,” Proceedings of the National Academy of Sciences, vol. 109, no. 24, pp. 9499–9504, 2012.

[59] T. J. Broering, K. A. Garrity et al., “Identification and characterization of broadly neutralizing human monoclonal antibodies directed against the E2 envelope glycoprotein of hepatitis C virus,” Virology, vol. 83, no. 23, pp. 12 473–12482, 2009.

[60] Z.-Y. Keck, T.-K. Li et al., “Analysis of a highly flexible conformational immunogenic domain a in hepatitis C virus E2,” Virology, vol. 79, no. 21, pp. 13199–13208, 2005.

[61] L. Kong, D. E. Lee et al., “Structural flexibility at a major conserved antibody target on hepatitis C virus E2 antigen,” Proceedings of the National Academy of Sciences, vol. 113, no. 45, pp. 12 768–12 773, 2016.

[62] M. Law, T. Maruyama et al., “Broadly neutralizing antibodies protect against hepatitis C virus quasispecies challenge,” Nature Medicine, vol. 14, no. 1, pp. 25–27, 2008.

[63] S. J. Merat, C. Bru et al., “Cross-genotype AR3-specific neutralizing antibodies confer long-term protection in injecting drug users after HCV clearance,” Hepatology, vol. 71, no. 1, p. 1424, 2019.

[64] V. Dahirel, K. Shekhar et al., “Coordinate linkage of HIV evolution reveals regions of immunological vulnerability,” Proceedings of the National Academy of Sciences, vol. 108, no. 28, pp. 11 530–11 535, 2011.

[65] A. A. Quadeer, R. H. Y. Louie et al., “Statistical linkage analysis of substitutions in patient-derived sequences of genotype 1a hepatitis C virus nonstructural protein 3 exposes targets for immunogen design,” Journal of Virology, vol. 88, no. 13, pp. 7628–7644, 2014.

[66] A. A. Quadeer, D. Morales-Jimenez, and M. R. McKay, “Co-evolution networks of HIV/HCV are modular with direct association to structure and function,” PLOS Computational Biology, vol. 14, no. 9, p. e1006409, 2018.

[67] S. F Ahmed, A. A. Quadeer et al., “Sub-dominant principal components inform new vaccine targets for HIV Gag,” Bioinformatics, vol. 35, no. 20, pp. 3884–3889, 2019.

[68] G. D. Gaiha, E. J. Rossin et al., “Structural topology defines protective CD8+ T cell epitopes in the HIV proteome,” Science, vol. 364, no. 6439, pp. 480–484, 2019.

[69] S. L. Chen and T. R. Morgan, “The natural history of hepatitis C virus (HCV) infection,” International Journal of Medical Sciences, pp. 47–52, 2006.

[70] S. B. Missiha, M. Ostrowski, and E. J. Heathcote, “Disease progression in chronic hepatitis C: Modifiable and nonmodifiable factors,” Gastroenterology, vol. 134, no. 6, pp. 1699–1714, 2008.

[71] Z. Yan and Y. Wang, “Viral and host factors associated with outcomes of hepatitis C virus infection,” Molecular Medicine Reports, vol. 15, no. 5, pp. 2909–2924, 2017.

[72] M. Rodríguez-Lopez, M. Ruiz et al., “Immunogenicity of variable regions of hepatitis C virus proteins: Selection and modification of peptide epitopes to assess hepatitis C virus genotypes by ELISA.” Journal of General Virology, vol. 80, no. 3, pp. 727–738, 1999.

[73] R. Vita, S. Mahajan et al., “The Immune Epitope Database (IEDB): 2018 update,” Nucleic Acids Research, vol. 47, no. D1, pp. D339–D343, 2018.

[74] P. J. Easterbrook, M. Smith et al., “Impact of HIV-1 viral subtype on disease progression and response to antiretroviral therapy,” Journal of the International AIDS Society, vol. 13, no. 1, p. 4, 2010.

[75] P. N. Amornkul, E. Karita et al., “Disease progression by infecting HIV-1 subtype in a seroconverter cohort in sub-saharan africa,” AIDS, vol. 27, no. 17, pp. 2775–2786, 2013.

[76] D. Ssemwanga, R. N. Nsubuga et al., “Effect of HIV-1 subtypes on disease progression in rural Uganda: A prospective clinical cohort study,” PLoS ONE, vol. 8, no. 8, pp. 1–7, 2013.

[77] N. Kiwanuka, A. Ssetaala et al., “High HIV-1 prevalence, risk behaviours, and willingness to participate in HIV vaccine trials in fishing communities on Lake Victoria, Uganda,” Journal of the International AIDS Society, vol. 16, no. 1, p. 18621, 2013.

[78] D. T. Claiborne, J. L. Prince et al., “Replicative fitness of transmitted HIV-1 drives acute immune activation, proviral load in memory CD4+ T cells, and disease progression,” Proceedings of the National Academy of Sciences, vol. 112, no. 12, pp. E1480–E1489 2015.

[79] J. R. Bailey, E. Barnes, and A. L. Cox, “Approaches, progress, and challenges to hepatitis C vaccine development,” Gastroenterology, vol. 156, no. 2, pp. 418–430, 2018.

[80] S. Muyldermans, “Nanobodies: Natural single-domain antibodies,” Annual Review of Biochemistry, vol. 82, no. 1, pp. 775–797, 2013.

[81] P. Sormanni, F. A. Aprile, and M. Vendruscolo, “Rational design of antibodies targeting specific epitopes within intrinsically disordered proteins,” Proceedings of the National Academy of Sciences, vol. 112, no. 32, pp. 9902–9907, 2015.

[82] I. Zimmermann, P. Egloff et al., “Synthetic single domain antibodies for the conformational trapping of membrane proteins,” eLife, vol. 7, p. e34317, 2018.

[83] K. Strimmer and A. V. Haeseler, “Genetic distances and nucleotide substitution models,” in The phylogenetic handbook: A practical approach to DNA and protein phylogeny, P. Lemey, M. Salemi, and A.-M. Vandamme, Eds. Cambridge University Press, 2009, pp. 112–113.

[84] C. Leys, C. Ley et al., “Detecting outliers: Do not use standard deviation around the mean, use absolute deviation around the median,” Journal of Experimental Social Psychology, vol. 49, no. 4, pp. 764–766, 2013.

[85] A. A. Quadeer, M. R. McKay et al., “MPF-BML: A standalone GUI-based package for maximum entropy model inference,” Bioinformatics, vol. 36, no. 7, pp. 2278–2279, 2019.

[86] M. Ekeberg, C. Lövkvist et al., “Improved contact prediction in proteins: Using pseudolikelihoods to infer potts models,” Physical Review E, vol. 87, no. 1, p. 012707, 2013.

[87] L. Kong, E. Giang et al., “Hepatitis C virus E2 envelope glycoprotein core structure,” Science, vol. 342, no. 6162, pp. 1090–1094, 2013.

[88] S. Miller, J. Janin et al., “Interior and surface of monomeric proteins,” Journal of Molecular Biology, vol. 196, no. 3, pp. 641–656, 1987.

[89] J. G. Jardine, D. Sok et al., “Minimally mutated HIV-1 broadly neutralizing antibodies to guide reductionist vaccine design,” PLoS Pathogens, vol. 12, no. 8, pp. 1–33, 2016.

## Supplementary References

[1] A. L. Ferguson, J. K. Mann et al., “Translating HIV sequences into quantitative fitness landscapes predicts viral vulnerabilities for rational immunogen design,” Immunity, vol. 38, no. 3, pp. 606–617, 2013.

[2] F. Morcos, A. Pagnani et al., “Direct-coupling analysis of residue coevolution captures native contacts across many protein families,” Proceedings of the National Academy of Sciences, vol. 108, no. 49, pp. E1293–E1301, 2011.

[3] R. Vita, S. Mahajan et al., “The Immune Epitope Database (IEDB): 2018 update,” Nucleic Acids Research, vol. 47, no. D1, pp. D339–D343, 2018.

[4] Z.-Y. Keck, C. Girard-Blanc et al., “Antibody response to hypervariable region 1 interferes with broadly neutralizing antibodies to hepatitis C virus,” Virology, vol. 90, no. 6, pp. 3112–3122, 2016.

[5] N. Kato, H. Sekiya et al., “Humoral immune response to hypervariable region 1 of the putative envelope glycoprotein (gp70) of hepatitis C virus.” Virology, vol. 67, no. 7, pp. 3923–3930, 1993.

[6] Z.-Y. Keck, O. Olson et al., “A point mutation leading to hepatitis C virus escape from neutralization by a monoclonal antibody to a conserved conformational epitope,” Virology, vol. 82, no. 12, pp. 6067–6072, 2008.

[7] Z.-Y. Keck, S. H. Li et al., “Mutations in hepatitis C virus E2 located outside the CD81 binding sites lead to escape from broadly neutralizing antibodies but compromise virus infectivity,” Virology, vol. 83, no. 12, pp. 6149–6160, 2009.

[8] T. J. Morin, T. J. Broering et al., “Human monoclonal antibody HCV1 effectively prevents and treats HCV infection in chimpanzees,” PLoS Pathogens, vol. 8, no. 8, p. e1002895, 2012.

[9] J. R. Bailey, L. N. Wasilewski et al., “Naturally selected hepatitis C virus polymorphisms confer broad neutralizing antibody resistance,” Journal of Clinical Investigation, vol. 125, no. 1, pp. 437–447, 2015.

[10] R. Velazquez-Moctezuma, A. Galli et al., “Hepatitis C virus escape studies of human antibody AR3A reveal a high barrier to resistance and novel insights on viral antibody evasion mechanisms,” Virology, vol. 93, no. 4, pp. e01 909–18, 2019.

